# Population dynamics of decision making in temperate bacteriophages

**DOI:** 10.1101/2020.03.18.996918

**Authors:** Moritz Lang, Maroš Pleška, Cǎlin C. Guet

## Abstract

Due to their ability to choose between lysis and lysogeny, temperate bacteriophages represent a classic model system to study the molecular basis of decision making. The coinfection of individual bacteria by multiple, genetically identical phages is known to alter the infection outcome and favor lysogeny over lytic development. However, it is not clear what role the ability of individual phages to sense and respond to coinfections plays in the phage-host infection dynamics at the population level. To address this question, we developed a full-stochastic model to capture the interaction dynamics between billions of bacteria and phages with single-cell and -phage resolution. While, at the level of individual bacteria, the probability of coinfections depends mainly on the phage concentration at the time of infection, the average number of coinfections at the population level is primarily determined by the relative growth rate of phage. Because the maximum attainable phage growth rate is constrained by basic life history parameters, the average number of coinfections has an upper bound of around two. However, for a broad range of conditions, the average number of coinfections stays well below this value. Consequently, we find that coinfections provide only very limited information to individual phages about the state of the infection at the population level. Nevertheless, this information can still provide a strong competitive advantage for phages that base fate decisions on the number of coinfections.

## Introduction

Phages are an integral part of the bacterial world and outnumber bacteria in many environments (1; 2; 3). Numerous theoretical and experimental studies examined the interaction dynamics between bacteria and phages at the population level (3; 4; 5; 6; 7; 8; 9; 10; 11; 12) (13; 14; 15; 16). However, the vast majority of these studies focused on obligatory lytic (also known as virulent) phages (4; 5; 6; 7; 8; 9; 10; 16), which reproduce exclusively horizontally at the expense of their hosts. In contrast, the interaction dynamics between bacteria and temperate phages (see below) were subject to only few studies (3; 11; 12; 13; 14; 15). While the molecular biology of several temperate phages is relatively well-understood (17; 18), many questions regarding their role in the ecology and evolution of bacteria remain open (19).

Temperate phages are defined by their ability to enter the cytoplasm of bacteria and coexist with them in a quasi-stable state called lysogeny (18; 20; 21). A typical infection by a temperate phage can lead to either: (i) lysis, accompanied by horizontal reproduction of the phage; or (ii) lysogeny, characterized by an indefinite number of vertical reproduction cycles together with the bacterial genome as a prophage (when referring to the phage) or a lysogen (when referring to both the bacterium and the phage). The molecular mechanisms that underlie the decision of the phage to either lyse or lysogenize the host bacterium have been studied in particular detail on the example of phage λ (18; 22). While probabilistic in nature, it has been shown that the decision-making process is, to a large extent, dependent on population-level parameters such as the bacterial growth phase (20), as well as on parameters that are masked at the population level, such as variability in cell size (23). Another important parameter influencing the lyse/lysogenize decision is the number of genetically identical phages coinfecting the same bacterium (24; 25). Based on experimental data for phage λ (24), it was estimated that the probability to be lysogenized can be as low as 1% for bacteria infected by a single phage, but close to 100% when coinfected by three or more phages (26), the exact values being dependent on the experimental conditions (25; 27; 28). Similar results were obtained for other temperate phages and bacterial hosts (29; 30), indicating that the ability of temperate phages to respond to multiple coinfections plays an important role in their ecology and could represent an evolved trait.

While the dependency of the fate decision on the number of genetically identical coinfecting phages is well understood (24; 25; 26; 27; 31; 32; 33; 34), little is known about the number of coinfections encountered by individual bacteria while the phage spreads in the bacterial population. Furthermore, the question of how the ability of temperate phages to sense and respond to multiple coinfections influences the phage-host interaction dynamics at the population level remains unanswered (34). In order to address these questions, we constructed a stochastic model of the interaction dynamics between billions of bacteria and temperate phages and analyzed how the probability that multiple phages coinfect the same bacterium changes as the phage spreads in the bacterial population. This stochastic model allows to precisely count the number of coinfections for every individual bacterium, and thus represents an unbiased approach where all the statistics on coinfections are directly determined by first-order principles of diffusion.

## Results

### Preliminaries

In our stochastic model (see Materials & Methods), we consider a well-mixed population of exponentially growing bacteria that is infected by an initially small number of phages (Fig. 1A). We assume that the environmental conditions allow the phage to take over the bacterial population in a few hours by means of successive horizontal reproduction cycles. We do not consider conditions in which such a fast spread of the phage cannot occur, e.g. due to concentrations of susceptible bacteria being too low (35; 16), nor conditions characterized by long term coexistence of phages and susceptible bacteria at quasi-steady concentrations (4; 6; 8; 12). We assume the interactions between individual bacteria and phages to follow a three-step process (Fig. 1B): (i) susceptible bacteria and phages freely diffuse in a well-mixed habitat and collide randomly, resulting in an infection; (ii) in the subsequent time interval, which we refer to as *fate decision period* Δ*t*_*f*_, infected bacteria randomly collide with other phages, leading to coinfections that can influence the infection outcome (Fig. 1B); (iii) after the end of the fate decision period Δ*t*_*f*_, the duration of which we estimated from recent single-cell experimental studies (see Materials & Methods and (27)), the infecting phages commit to one of two possible pathways leading to either the integration of the phage DNA into the host genome (lysogeny) or to the production of progeny phages, accompanied by bursting of the host bacterium (lysis). The time interval from the first phage adsorption to lysis, which also includes the fate decision period Δ*t*_*f*_, is referred to as the *latent period* Δ*t*_*l*_ (Fig. 1B).

**Fig. 1.**
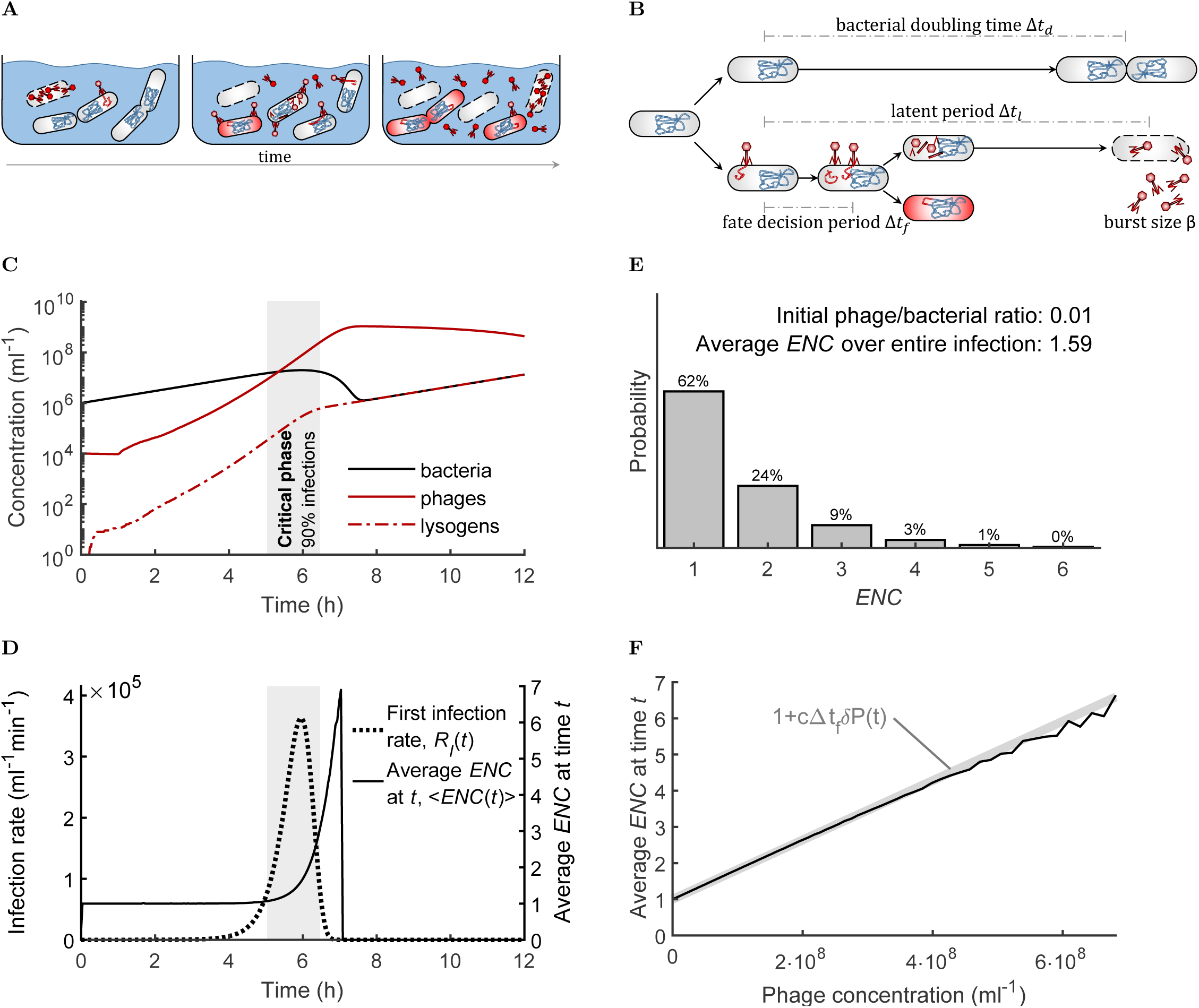
Dynamics of a bacterial population infected by temperate phages. (A) A growing population of susceptible bacteria (gray cells) is infected by an initially small population of temperate phages (red particles) in a well-mixed habitat. The phages can either reproduce inside the infected bacteria leading to lysis, or integrate into the genome and establish lysogeny (red cells). Lysogens still divide and adsorb phages, but are immune to subsequent infections. (B) The time from infection to lysis is denoted as the latent period Δ*t*_*l*_, and typically has a duration similar in magnitude to the bacterial doubling time Δ*t*_*d*_ (37; 38). Together with the burst size *β*, the latent period Δ*t*_*l*_ determines the maximal relative growth rate of the phage population. The fate decision period Δ*t*_*f*_ corresponds to the time between the first infection and the time after which the influence of subsequent coinfections on the fate decision can be neglected. (C) Simulation of a bacterial population (black solid curve) infected by a temperate phage (red solid curve). The phages quickly spread and turn a fraction of the bacterial population into lysogens (red dash-dotted curve). (D) The rate *R*_*I*_(*t*) at which susceptible bacteria are infected by their first phage (dotted curve, left axis) dynamically changes during the infection dynamics. In contrast, the average *ENC* of a bacterium infected at time *t*, ⟨*ENC*(*t*)⟩ (solid curve, right axis), remains close to 1 for most of the time and only peaks sharply towards the end of the infection dynamics when the rate of new infections is already quickly decreasing. The gray area depicts the critical phase, i.e. the time interval during which 90% of all first infections occur. (E) Distribution of the *ENC* encountered by susceptible bacteria during the infection dynamics. Note that the *ENC*s are not Poisson distributed since the number of uninfected bacteria at the end of the simulation was zero. (F) The average *ENC* of a bacterium infected at time *t* increases approximately linearly with the free phage concentration at the same time *t* (black curve, corresponds to simulation shown in (C)), and is well approximated by Eq. 1 (light gray line, corresponds to *c*(*t*) = 1.2). All simulations were performed for a total volume of 1*ml*, and with initial bacterial and phage concentrations of 10^6^*ml*^−1^ and 10^4^*ml*^−1^, respectively. The probability of lysogenization was fixed to 2%, corresponding to the *ENC*-insensitive phage variant. For the corresponding dynamics of the *ENC*-sensitive phage variant, see Supplementary Figure 2.

We refer to the total (integer) number of phage coinfections that occur during the fate decision period of an individual bacterium as the *effective number of coinfections* (*ENC*). Thus, we only count towards the ENC those coinfections that occur early enough to significantly influence the fate decision, even though we assume that phages can still adsorb to an infected bacterium after its fate decision period has ended (as well as to lysogens). Note, that the *ENC* is a variable defined at the level of individual bacteria and phages, and as such, has to be distinguished from the multiplicity of infection (*MOI*) which typically denotes the phage/bacteria ratio at the time of inoculation (Box 1). Importantly, the *MOI* does in general not quantify coinfections, except under particular experimental conditions specifically designed to circumvent several effects arising from the phage-host interaction dynamics at the population level (see Box 1, Supplementary Figure 1, and (24)).

#### Box 1.

**Quantifying the number of coinfections by multiple phages**

**Box 1.**
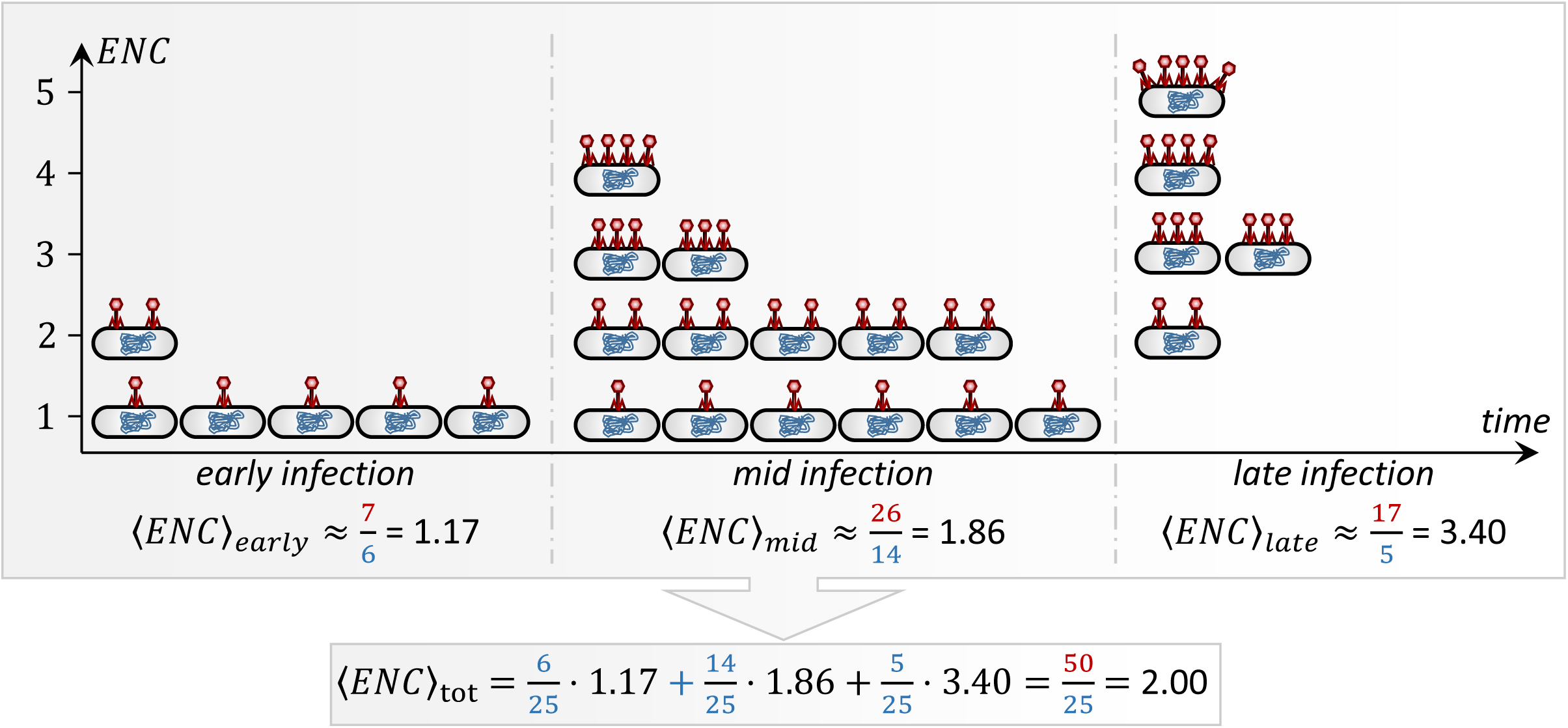

The effective number of coinfections (*ENC*) denotes the number of coinfections of a given bacterium by multiple phages of the same kind which occur during the fate decision period and can thus influence the infection outcome. The *ENC* is defined at the level of individual bacteria and has to be distinguished from the multiplicity of infection (*MOI*), which typically correspond to the phage/bacteria ratio at the time of inoculation (5 p. 40) and is thus defined at the level of populations of phages and bacteria (for a discussion on alternative definitions of the *MOI*, see (56)). The *ENC* is conceptually similar to a quantity which is sometimes referred to as the “cellular *MOI*”, the total number of coinfections of an individual bacterium (34; 27), typically assuming equal arrival times of all coinfecting phages. However, unlike the “cellular *MOI*”, the *ENC* only takes into account those phage coinfections that occurred during the fate decision period, i.e. before the cellular decision to lyse or lysogenize was already made.

The average *ENC* of a bacterium infected by its first phage at time *t*, denoted as ⟨*ENC*(*t*)⟩, can be approximated by the mean value of the *ENCs* of all bacteria infected during a small time interval around *t*, and increases approximately linearly with the phage concentration (see main text). When calculating the average *ENC* encountered by bacteria over the entire infection dynamics (denoted as ⟨*ENC*⟩_*tot*_), the ⟨*ENC*(*t*)⟩ has to be weighted by the rate *R*_*I*_(*t*) = *δB*^−^(*t*)*P*(t) of infections of susceptible bacteria by their first phage (see schematic):

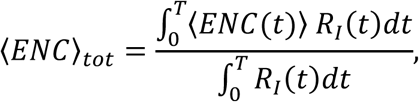

where *B*^−^(*t*) and *P*(*t*) denote the concentrations of uninfected bacteria and phages, respectively, and *δ* denotes the adsorption rate constant. Importantly, the ⟨*ENC*⟩_*tot*_ only takes into account those bacteria infected by at least one phage and is thus always greater or equal to one. Note that the ⟨*ENC*⟩_*tot*_ and the fraction of bacteria which got infected during an experiment are in general independent and should thus be reported separately (compare (56)).

Due to the nonlinear dynamics of the rate of new infections, the *ENC*s encountered by bacteria during the entire infection dynamics are in general not Poisson distributed. Furthermore, the ⟨*ENC*⟩_*tot*_ is in general neither given nor well approximated by the phage/bacterial ratio at the time of inoculation (i.e. the *MOI*) or at any other time, and can even be anti-correlated to the *MOI* (see main text). Since the average “cellular *MOI*” would have the same value as the ⟨*ENC*⟩_*tot*_ for the (unrealistic) case of equal fate decision and latent periods, this implies that also the average “cellular *MOI*” is in general not correlated to the *MOI*.

Throughout the manuscript, we distinguish between *ENC*-sensitive and -insensitive phage “variants”. For the *ENC*-sensitive variant, we assume that the probability of lysogenization shows a strong dependency on the number of coinfections as experimentally observed for phage λ (0.38%, 69.60% and 98.86% for an *ENC* of one, two, and more than two, respectively; see (26)). For the (hypothetical) *ENC*-insensitive variant, we in contrast assume that the probability of lysogenization is constant (2% if not noted otherwise). In the first part of the manuscript, we analyze general properties of the infection dynamics which hold for both variants, i.e. independently of whether we assume an *ENC*-dependent probability of lysogenization or not. This serves as the foundation for the second part of the manuscript, where we analyze the dynamic differences between the two variants and thus the role of the phages’ ability to sense and respond to multiple coinfections in the phage-host interaction dynamics.

### Infection dynamics and coinfections by multiple phages

The stochastic simulations of our model show infection dynamics similar to recent experimental studies (11). Initially, the phage concentration is too low to have any significant effect on the bacterial population, and the bacterial concentration grows exponentially due to cell division (Fig. 1C for the *ENC*-insensitive phage variant and Supplementary Figure 2A for the *ENC*-sensitive one). As the phage concentration increases in subsequent cycles of infections, also the rate at which susceptible bacteria are infected by their first phage steadily increases (Fig. 1C&D and Supplementary Figure 2A&B). When this rate becomes sufficiently high (see below), the phage takes over the bacterial population (Fig. 1C&D and Supplementary Figure 2A&B). Due to the fast growth of the phage, the bulk of infections thereby occurs during a relatively short *critical phase* of the infection dynamics, which we define as the narrowest time interval during which 90% of all susceptible bacteria are infected by their first phage (Fig. 1C&D and Supplementary Figure 2A&B). Since the critical phase has a similar duration as the latent period, it comprises the last horizontal reproduction cycle of the phage before the vast majority of susceptible bacteria is already infected (Fig. 1C&D and Supplementary Figure 2A&B). After the critical phase, whose end is characterized by the start of a sharp decline in the total bacterial concentration (Fig. 1C and Supplementary Figure 2A), nearly all bacteria are either lysogenic or in the process to lyse/lysogenize.

**Fig. 2.**
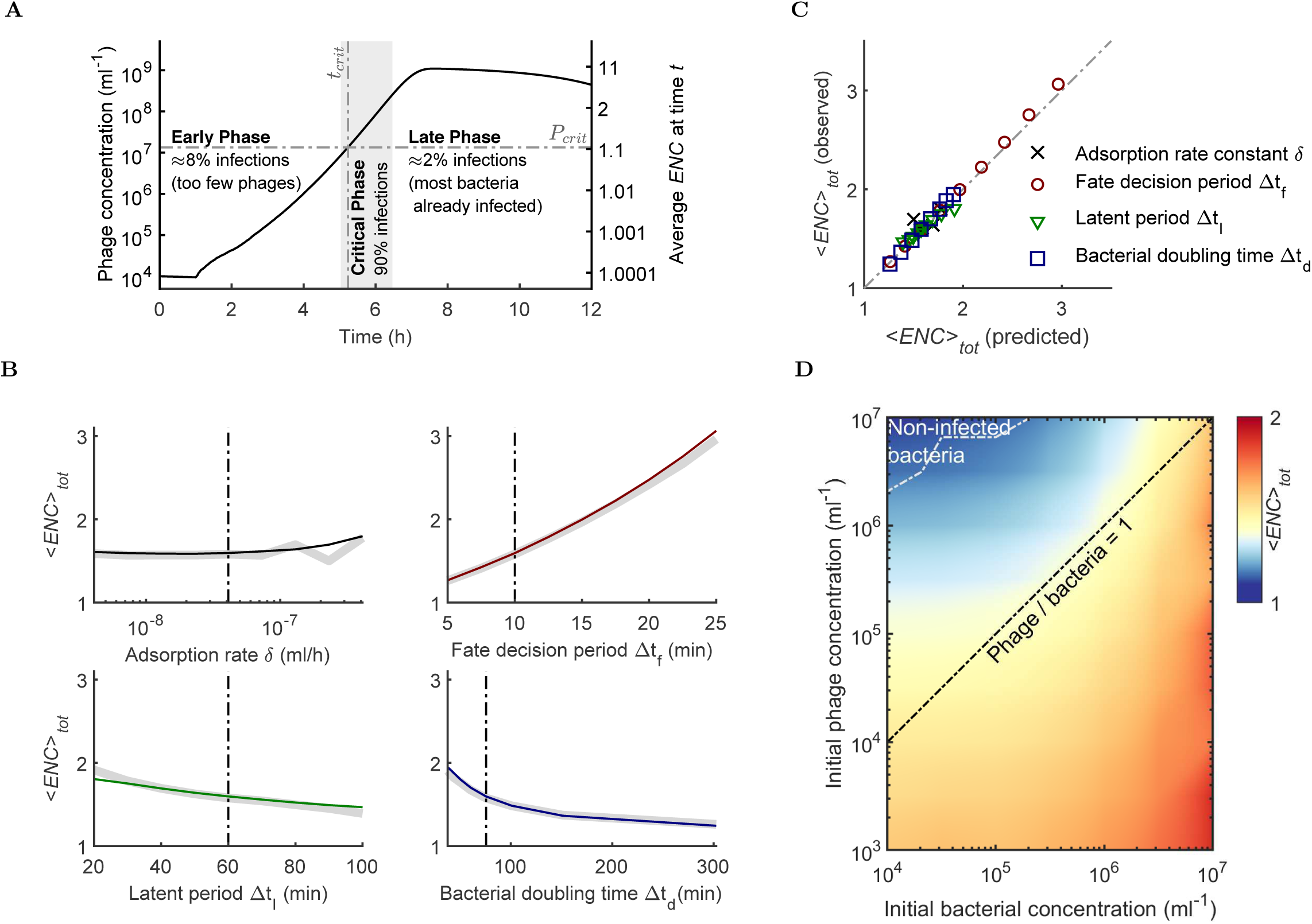
Dependency of the *ENC* on key environmental parameters. (A) By definition, 90% of all susceptible bacteria are infected during the relatively short critical phase (gray area) of the infection dynamics, which starts approximately at the time *t*_*crit*_ when the phages reach the critical concentration threshold *P*_*crit*_ (see main text). The black curve represents the same phage dynamics *P*(*t*) over time *t* as in Fig. 1C (left axis), as well as the corresponding average *ENC* of bacteria infected at the same time, ⟨*ENC*(*t*)⟩ (right axis). (B) Relationship between the average *ENC* over the entire infection dynamics, ⟨*ENC*⟩_*tot*_, and key parameters of our model (colored curves). Vertical dash-dotted lines represent the respective reference values of the parameters (Table 1), and light gray curves the prediction of Eq. 2. (C) Comparison between the ⟨*ENC*⟩_*tot*_ obtained from the simulations in and the value directly calculated via our analytic formula (compare B). (D) The ⟨*ENC*⟩_*tot*_ as a function of the initial bacterial (10^4^ – 10^7^*ml*^−1^) and phage (10^3^ – 10^7^*ml*^−1^) concentrations at the time of inoculation. Red colors represent high values of the ⟨*ENC*⟩_*tot*_, and blue colors represent low values. The area inside the white dash-dotted curve indicates a zone of high initial phage and low initial bacterial concentrations, for which a small fraction of susceptible bacteria remained uninfected at the end of the simulation (*T* = 16*h*). The black dash-dotted line corresponds to an initial phage/bacteria ratio of one. Note, that the phage/bacteria ratio at the time of inoculation (i.e. the *MOI*) increases towards the top-left of the plot, while the ⟨*ENC*⟩_*tot*_ decreases in this direction. All simulations were performed for a total volume of 1*ml*. Parameter sensitivities were determined for initial bacterial and phage concentrations of 10^6^*ml*^−1^ and 10^4^*ml*^−1^, respectively. The probability of lysogenization was fixed to 2%, corresponding to the *ENC*-insensitive phage variant. For the corresponding analysis of the *ENC*-sensitive phage variant, see Supplementary Figure 3.

Based on the stochastic simulations described above, we extracted the *ENC*s by directly counting the exact number of phage coinfections that occurred during the fate decision period of each infected bacterium. Based on the *ENC*s of all infected bacteria, we calculated the distribution of the *ENC*s and the average *ENC* over the whole infection dynamics, denoted ⟨*ENC*⟩_*tot*_. From the simulations of both phage variants, we found a value of ⟨*ENC*⟩_*tot*_ of only around 1.6, and that the majority of bacterial fate decisions (62%) was based on a single phage infection (Fig. 1E and Supplementary Figure 2C). To analyze how the probability for multiple coinfections changes as the phages spread in the bacterial population, we further estimated the average *ENC* for bacteria infected at a given time *t* during the infection dynamics, denoted ⟨*ENC*(*t*)⟩. For both phage variants, the ⟨*ENC*(*t*)⟩ remained close to one throughout most of the simulation, and only peaked sharply near the end of the infection dynamics when the rate of new infections was already sharply decreasing (Fig. 1D and Supplementary Figure 2B). These results imply that coinfections by multiple phages only become likely for bacteria infected late during the infection dynamics, while during most of the infection dynamics nearly every fate decision is based on a single phage infection.

**Table 1.**
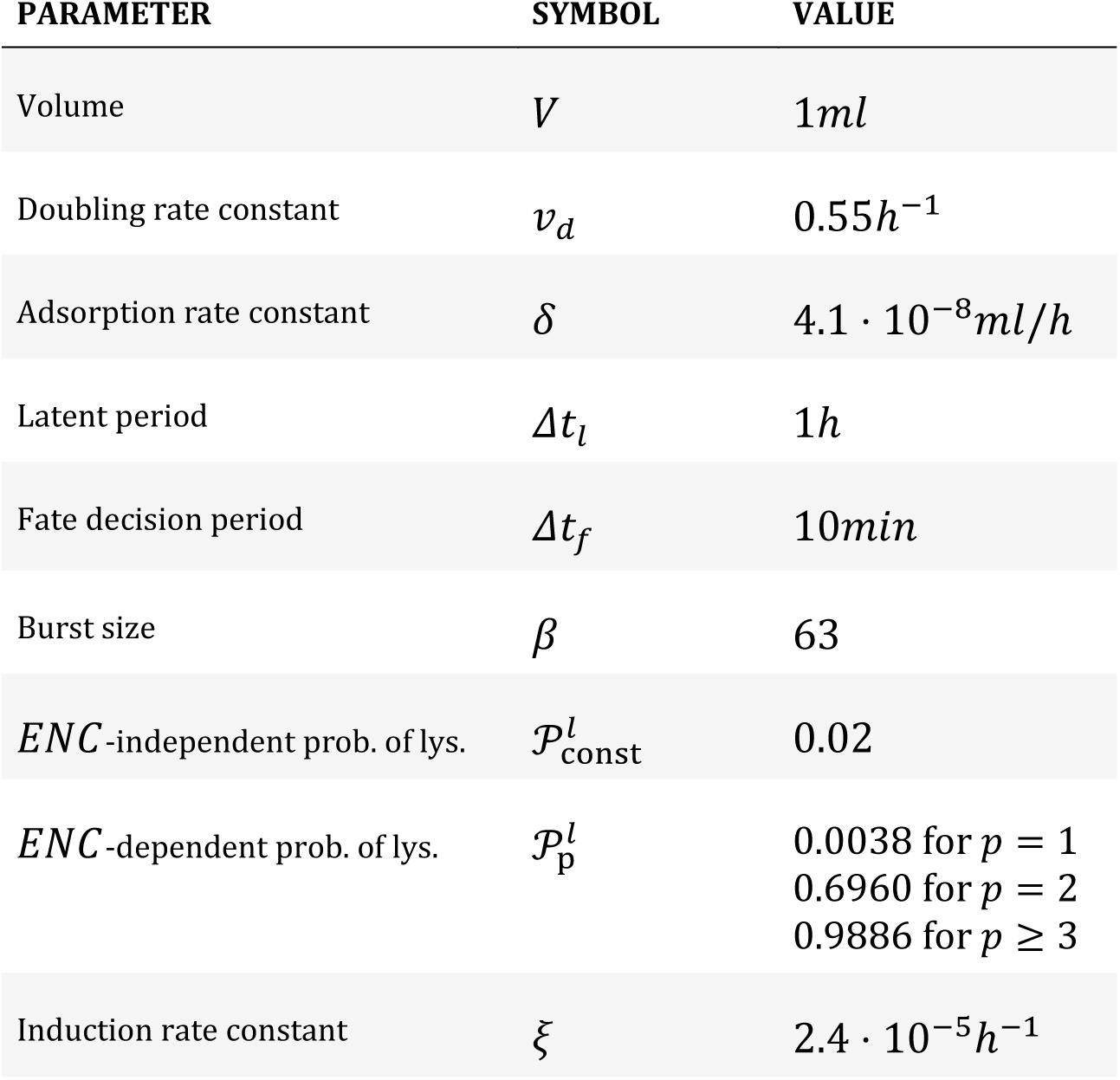
Parameters for our stochastic model of phage-host population dynamics.

| PARAMETER | SYMBOL | VALUE |
| --- | --- | --- |
| Volume | $V$ | $1ml$ |
| Doubling rate constant | $v_d$ | $0.55h^{-1}$ |
| Adsorption rate constant | $\delta$ | $4.1 \cdot 10^{-8}ml/h$ |
| Latent period | $\Delta t_l$ | $1h$ |
| Fate decision period | $\Delta t_f$ | $10min$ |
| Burst size | $\beta$ | 63 |
| *ENC*-independent prob. of lys. | $\mathcal{P}_{const}^l$ | 0.02 |
| *ENC*-dependent prob. of lys. | $\mathcal{P}_p^l$ | 0.0038 for $p = 1$<br>0.6960 for $p = 2$<br>0.9886 for $p \geq 3$ |
| Induction rate constant | $\xi$ | $2.4 \cdot 10^{-5}h^{-1}$ |

The number of phage coinfections per bacterium is often assumed to be approximately Poisson distributed with a mean given by the ratio between phage and bacterial concentrations at the time of inoculation, i.e. the *MOI* (5 p. 40ff; 24). However, this relationship is only valid for specific infection scenarios in which: (i) phage replication by lysing bacteria can be neglected, (ii) bacterial concentrations remain unchanged, and (iii) sufficient time is available for the majority of the inoculated phages to adsorb to bacteria (36) (see Supplementary Figure 1, and (24) for corresponding experimental procedures). Importantly, none of these requirements are satisfied in infection scenarios in which phages and bacteria interact over longer periods of time, characterized by dynamic changes in their respective concentrations, e.g. due to the bursting of lysing bacteria and the accompanied release of phage progeny. As a consequence, the distribution of the *ENC*s is not Poissonian and the average *ENC* over the entire infection dynamics, ⟨*ENC*⟩_*tot*_, is neither given nor well approximated by the phage/bacteria ratio at the time of inoculation or at any other time (Fig. 1E and and Supplementary Figure 2C). Instead, since the probability per unit of time that a bacterium adsorbs a phage only depends on the phage concentration at the respective time (5 p. 49ff), the average *ENC* of a bacterium infected at a given time *t*, ⟨*ENC*(*t*)⟩, increases approximately linearly with the phage concentration *P*(*t*) (Fig. 1F and Supplementary Figure 2D):

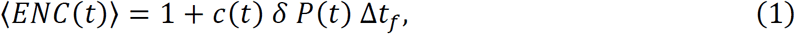

where Δ*t*_*f*_ denotes the fate decision period, *δ* the adsorption rate constant, and *c*(*t*) a factor taking into account the increase of phage concentration during the fate decision period. The factor 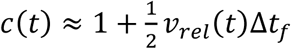 depends on the relative growth rate 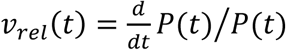 the phage (growth in percent of the current concentration), and typically has a value between 1 (slow phage growth) and 1.4 (fast phage growth; see Supplementary Text). In Figure 1F and Supplementary Figure 2D, we show that the results obtained from Eq. 1 agree with the values of the ⟨*ENC*(*t*)⟩ directly obtained from our stochastic simulations.

### The average number of coinfections and its determinants

To better understand how the number of coinfections depends on environmental conditions, we analyzed how the average *ENC* over the entire infection dynamics, ⟨*ENC*⟩_*tot*_, changes in response to variations of key parameters of our model. Mathematically, the ⟨*ENC*⟩_*tot*_ can be calculated as the weighted mean of the ⟨*ENC*(*t*)⟩, where the weight factor is proportional to the rate at which susceptible bacteria are infected by their first phage (Fig. 1D, see Box 1). Because 90% of all infections occur during the critical phase (Fig. 1C&D), due to this weighting, the ⟨*ENC*⟩_*tot*_ nearly exclusively depends on the population dynamics during the critical phase.

The time interval of the critical phase depends on the full infection dynamics and its precise mathematical analysis is thus challenging. However, the onset of the critical phase can be well approximated by the time *t*_*crit*_ when the concentration of uninfected bacteria stops to increase and starts to decrease, i.e. when the rate of decrease *δB*^−^(*t*)*P*(*t*) of uninfected bacteria *B*^−^(*t*) due to infections by phages *P*(*t*) balances the rate of increase *v*_*d*_*B*^−^(*t*) due to cell division, with *δ* being the adsorption rate constant and *v*_*d*_ the bacterial doubling rate constant (Supplementary Figure 4). When we set both rates equal, *δB*^−^(*t*)*P*(*t*) = *v*_*d*_*B*^−^(*t*), we find that *t*_*crit*_ is completely determined by the phage dynamics, and that the onset of the critical phase occurs approximately at the time when the phage concentration reaches the critical threshold 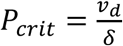 (referred to as the viral inundation threshold in the context of phage therapy (16)).

**Fig. 3.**
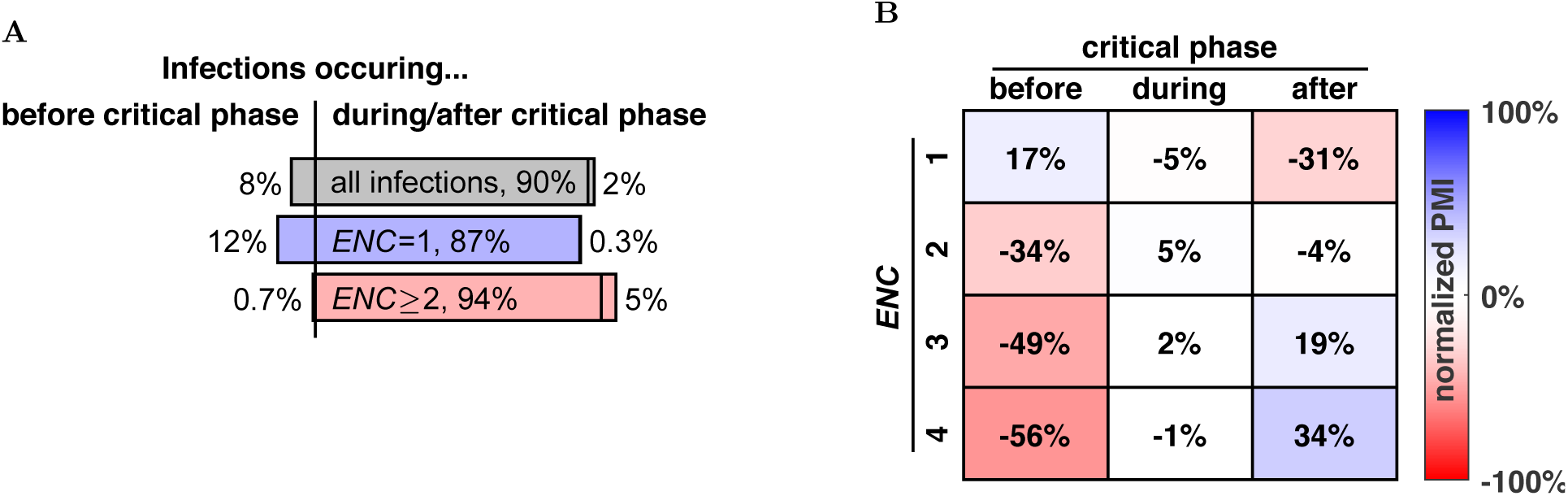
Information conveyed by the *ENC* to individual phages. (A) Fraction of first phage infections occurring before (left part of bar), during (central part) and after (right part) the critical phase when considering all infections (top), when only considering those infections with an *ENC* of one (middle), or greater than one (bottom). (B) Type of information conveyed by the *ENC* to individual phages, quantified by the normalized pointwise mutual information (*nPMI*) between the *ENC* and the phase of the infection. The *nPMI* is a measure of relative co-occurrence, with high positive values indicating a significantly increased, and high negative values a significantly decreased probability that a given *ENC* occurs before, during or after the critical phase (as compared to independence). The *nPMI*s with high negative values for multiple coinfections occuring before the critical phase indicate that the *ENC* mainly provides information to bacteria coinfected by multiple phages, in the form that their already rather low probability to belong to the initial phases of the infection is further decreased (compare A).

**Fig. 4.**
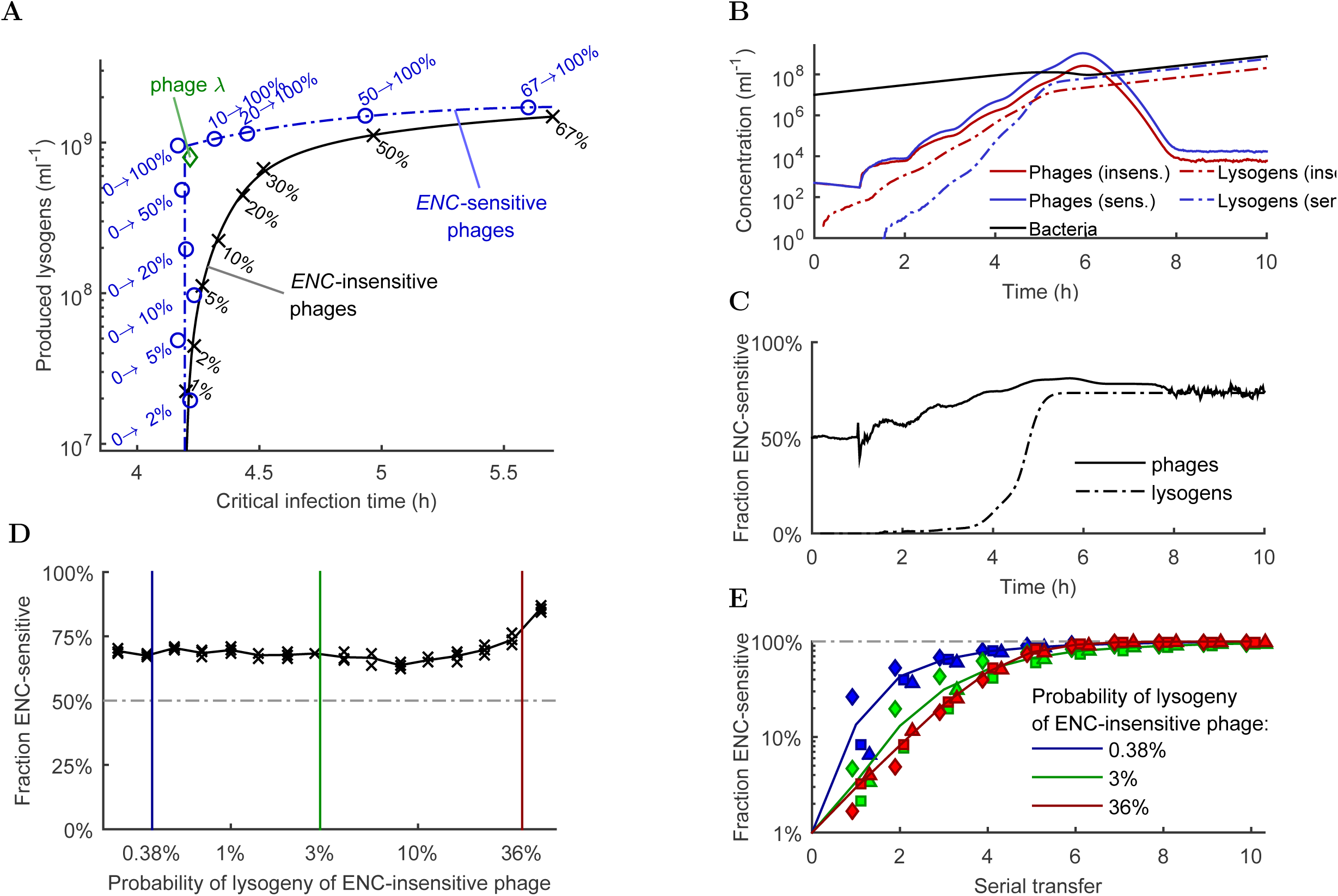
Attenuation of the trade-off between phage growth and lysogen yield by an *ENC*-dependent fate decision. (A) Relationship between final lysogen concentration and critical infection time for *ENC*-insensitive phages with different constant probabilities of lysogenization (black curve), and for an *ENC*-sensitive phages with a probability of lysogenization X → Y (blue curve), whereby X denotes the probability when coinfected by a single phage, and Y the probability when coinfected by two or more phages. The green diamond represents the experimentally observed probabilities of lysogenization for phage λ (0.38%, 69.60% and 98.86% for an *ENC* of one, two and more than two, respectively; see (26)). (B) Competition dynamics between *ENC*-sensitive (blue solid) and -insensitive (red solid) phages coinfecting a bacterial population with equal concentrations at the start of the simulation. The black curve corresponds to the total bacterial concentration, and the dashed blue and red curves to lysogens carrying the *ENC*-sensitive and -insensitive prophages, respectively. The probability of lysogenization for the *ENC*-sensitive phage was set to the values of phage λ (see A), and for the *ENC*-insensitive phages it was set to 36%. (c) Fraction of *ENC*-sensitive phages in the total phage population (black solid), and fraction of lysogens carrying the *ENC*-sensitive prophage in the total lysogen population (black dash-dotted) for the simulation shown in (B). (D) Final fraction of *ENC*-sensitive phages for competition experiments starting with equal numbers of *ENC*-sensitive and - insensitive phages (compare B&C) as a function of the constant probability of lysogenization of the *ENC*-insensitive phage (three independent simulations each). Vertical colored lines indicate the values of the probability of lysogenization corresponding to the ones used in (E). (E) Fraction of *ENC*-sensitive phages in serial transfer simulations. In the first transfer, the initial fraction of *ENC*-sensitive phages was set to 1%, and in all subsequent transfers to the respective final fraction of the previous transfer. The constant probability of lysogenization for the *ENC*-insensitive phage was set to 0.38% (blue), 3% (green) and 36% (red), respectively (three independent simulations each). All simulations were performed for an initial bacterial concentration of 10^7^*ml*^−1^ and an initial total phage concentration (*ENC*-sensitive & -insensitive) of 10^3^*ml*^−1^ at *t* = 0.

The phage dynamics are thus not only the main determinant of the average *ENC* of a bacterium infected at time *t*, ⟨*ENC*(*t*)⟩ (see previous section), but also of the onset of the critical phase. This allows us to directly approximate the average *ENC* at the onset of the critical phase by 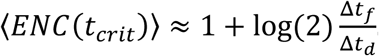. Since the fate decision period Δ*t*_*f*_ constitutes only a small fraction of the latent period Δ*t*_*l*_, and since the latent period typically has a duration similar to the bacterial doubling time 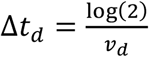 (37; 38) (Fig. 1B), it follows that the average *ENC* at the onset of the critical phase has to be rather small. For example, in the simulation shown in Fig. 1C&D, ⟨*ENC*(*t*_*crit*_)⟩ was 1.1 (Fig. 2A), implying that most fate decisions were still based on a single phage infection at the onset of the critical phase. Since the critical phase has a similar duration as the latent period, this also implies that the phage dynamics during the critical phase, and thus the average *ENC* over the entire infection dynamics, ⟨*ENC*⟩_*tot*_, are predominantly determined by the fate decisions of bacteria infected by a single phage. For the *ENC*-sensitive and -insensitive phage variants having both low absolute probabilities of lysogenization for infections by single phages, we indeed found nearly identical *ENC* distributions over the entire infection dynamics (Fig. 1E and Supplementary Figure 2C),

Since coinfections are still relatively rare at the onset of the critical phase, a high ⟨*ENC*⟩_*tot*_ can only be obtained if the phage replicates quickly and thus reaches high concentrations supporting large numbers of coinfections before the end of the critical phase when most bacteria are already infected. In agreement with this consideration, we derive in the Supplementary Information that the ⟨*ENC*⟩_*tot*_ increases approximately linearly with the product of the relative growth rate *v*_*rel*_(*t*_*crit*_) of the phage population at the onset *t*_*crit*_ of the critical phase, and the length Δ*t*_*f*_ of the fate decision period:

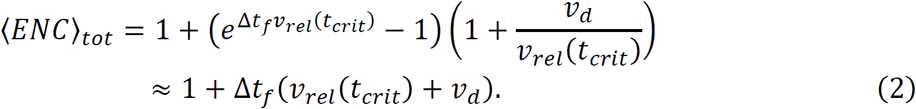

The relative phage growth rate *v*_*rel*_(*t*_*crit*_) and the bacterial doubling rate constant *v*_*d*_ can be directly estimated from the phage-host population dynamics: in a logarithmic plot of the phage and bacterial concentrations over time (Fig. 1C), *v*_*rel*_(*t*_*crit*_) and *v*_*d*_ correspond to the slope of the phage and bacterial concentrations shortly before the bacterial population dynamics become significantly influenced by the phage. By this method, we estimated a relative phage growth rate of *v*_*rel*_(*t*_*crit*_) = 2.29 *h*^−1^ and a bacterial doubling rate constant of *v*_*d*_ ≈ 0.55 *h*^−1^ for the infection dynamics shown in Fig. 1C, for which our formula predicts a value of ⟨*ENC*⟩_*tot*_ ≈ 1.58. This prediction is in good agreement with the value of 1.59 directly obtained from our stochastic simulations (Fig. 1E).

As a direct consequence of Eq. 2, every factor that leads to an increase of the phage growth rate will also lead to an increase of ⟨*ENC*⟩_*tot*_. In agreement with this theoretical prediction, we found in the simulations of our stochastic model that ⟨*ENC*⟩_*tot*_ increases for higher adsorption rate constants *δ* and shorter latent periods Δ*t*_*l*_ (Fig. 2B for the *ENC*-insensitive variant, Supplementary Figure 3B for the *ENC*-sensitive one). Because phages spread faster at higher bacterial concentrations, ⟨*ENC*⟩_*tot*_ increases with higher bacterial concentrations at the onset of the critical phase. Consequently, ⟨*ENC*⟩_*tot*_ increases both, if the initial bacterial concentration is higher (Fig. 2D and Supplementary Figure 3D) or if the bacterial concentration grows faster and thus reaches higher values during the critical phase (Fig. 2B and Supplementary Figure 3B). Similarly, a lower initial phage concentration leads to a later onset of the critical phase, which gives the bacterial population additional time to increase its concentration, leading to a higher relative phage growth rate during the critical phase and thus to a higher ⟨*ENC*⟩_*tot*_ (Fig. 2D and Supplementary Figure 3D). Finally, ⟨*ENC*⟩_*tot*_ also increases with prolonged durations of the fate decision period (Fig. 2B and Supplementary Figure 3B), as predicted by Eq. 2.

When we plotted the values of the ⟨*ENC*⟩_*tot*_ obtained from all simulations discussed above as a function of the corresponding predictions based on Eq. 2, all data points collapsed onto the main diagonal (Fig. 2C and Supplementary Figure 3C). In contrast, we found no clear correlation between ⟨*ENC*⟩_*tot*_ and the peak phage concentration (Supplementary Figure 5). Finally, we note that the results above also imply that ⟨*ENC*⟩_*tot*_ is in general anti-correlated to the initial phage/bacterial ratio (i.e. the *MOI*) under environmental conditions allowing for fast spread of phage in a susceptible bacterial population (Fig. 2D, Supplementary Figure 3D and Box 1).

### Limits of sensing

While phages can spread fast under favorable conditions characterized by high concentrations of susceptible bacteria, they cannot spread arbitrarily fast. The upper limit max(v_rel_) = log(*β*)/Δ*t*_*l*_ of the relative growth rate of the phage population is attained under very high concentrations of susceptible bacteria, i.e. when every released phage nearly immediately finds a bacterium to infect, which then results in the release of *β* (burst size) progeny phages after the latent period Δ*t*_*l*_. By inserting this upper limit into the equation for ⟨*ENC*⟩_*tot*_ (Eq. 2), we obtain an upper bound for the maximally attainable average *ENC* over the entire infection dynamics:

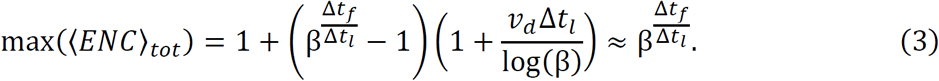

With a burst size *β* = 63, a latent period Δ*t*_*l*_ = 60*min* and a fate decision period Δ*t*_*f*_ = 10*min* (Table 1), we obtain a maximal attainable average *ENC* over the entire infection dynamics of only 2.1. The practically attainable values of the ⟨*ENC*⟩_*tot*_, however, usually stay well below this physical upper bound (Fig. 2A-D). Indeed, we could only observe values for ⟨*ENC*⟩_*tot*_ of 2 or higher if we significantly modified one of the three parameters (*β*, Δ*t*_*l*_, or Δ*t*_*f*_) the upper bound itself depends on (Eq. 3), e.g. when we assumed very long fate decision periods lasting for more than a quarter of the respective latent period (Fig. 2B).

To better understand the implications of this upper bound (Eq. 3) for the ability of individual phages to obtain information about the phase of the infection at the population level, we asked how the average *ENC* of those infections that occur before the critical phase, ⟨*ENC*⟩_*before*_, differs as compared to those infections that occur during the critical phase, ⟨*ENC*⟩_*crit*_. Since, before the critical phase, nearly all fate decisions are based on a single phage infection (see previous section), it directly follows that ⟨*ENC*⟩_*before*_ ≈ 1. On the other hand, since 90% of all infections occur during the critical phase, the average *ENC* during the critical phase is nearly identical to the average *ENC* over the full infection dynamics, i.e. ⟨*ENC*⟩_*crit*_ ≈ ⟨*ENC*⟩_*tot*_. The upper bound for ⟨*ENC*⟩_*tot*_ (Eq. 3) thus implies that even during the critical phase of the infection dynamics, which comprises the last horizontal reproduction cycle of the phage, a large fraction of bacterial fate decisions— typically the majority—is still based on infections by a single phage (Fig. 1E).

As a consequence of the high prevalence of infections with *ENC* = 1, both before and during the critical phase, coinfections can only convey very limited information to individual phages about the phase of the infection at the population level. For the infection dynamics shown in Fig. 1C, the mutual information between the phase of the infection (early, critical or late) and the *ENC* (taking integer values that represent numbers of coinfections) for example only amounted to *I(Phase; ENC*) = 0.08*bit*, significantly less than the 1*bit* necessary to conclusively answer a yes/no question. This indicates that the number of coinfections is not a sufficient environmental cue for individual phages to identify the phase of the infection at the population level. The quantitative lack of information conveyed by *ENC* has to be seen in comparison to the high prior probability of 90% that each individual infection occurs during the critical phase. The very limited information conveyed by *ENC* cannot significantly affect this prior probability, such that all infections, even those with an *ENC* of one, most likely occur shortly before the majority of susceptible bacteria is already infected. For the infection dynamics in Fig. 1C, for example, as much as 87% of infections with an *ENC* of one occur during the critical phase, only slightly less than the 90% of all infections and the 94% of infections with an *ENC* of two or more (Fig. 3A).

That the information conveyed by *ENC* to individual phages is not sufficient to discriminate between the different phases of the infection dynamics (and thus between the last and all previous reproduction cycles of the phage) however does not imply that sensing the *ENC* cannot lead to more informed fate decisions. When we analyzed the *type* of information transmitted by the *ENC* (as opposed to its mere *quantity*), we found that the *ENC* provides nearly no information concerning the role of the critical phase as the most likely time interval during which bacteria are infected, but rather informs the individual phage about their residual probabilities to belong to the phases before or after the critical phase (Fig. 3B). For the infection dynamics in Fig. 1C, an *ENC* of two or more, for example, provides the relative information that the already small prior probability that the infection occurred before the critical phase is even tenfold decreased, while not significantly increasing the already high prior probability of the critical phase itself (Fig. 3A). This information about the relative probabilities to belong to the early or late phases of the infection dynamics might still be utilized by phages basing their fate decision on the *ENC*, even though the information conveyed by *ENC* does not allow them to discriminate between the different phases of the infection dynamics in absolute terms.

### Tradeoff between phage growth and lysogenized bacteria

In stochastic simulations of our model for the *ENC*-insensitive phage variant having a constant probability of lysogenization, we found that the concentration of lysogens at the end of the infection dynamics monotonically increased with increasing probabilities of lysogenization (Fig. 4A). On the other hand, an increase in the probability of lysogenization also resulted in a later onset *t*_*crit*_ of the critical phase, i.e. increased the time required for the phage to take over the bacterial population (Fig. 4A). This increase of *t*_*crit*_ can be directly explained by the fact that phages which lysogenize infected bacteria do not immediately contribute to the production of new phage progeny, resulting in a lower growth rate of the phage population. These results indicate a trade-off between the rate at which temperate phages can spread in susceptible bacterial populations and the fraction of bacteria that they can lysogenize.

In principle, this trade-off could be circumvented if phages infecting bacteria during the early phase of the infection dynamics favored lysis while late infecting phages favored lysogeny, thus allowing for a quick spread of the phage while maximizing the fraction of lysogenized bacteria. However, as described in the previous section, the information conveyed by the *ENC* does not allow individual phages to discriminate between the phases of the infection dynamics (Fig. 3A). Because a strong bias towards lysis during the initial infection phase is nevertheless essential for a quick take-over of the bacterial population, this leads to the seemingly paradoxical situation where *some* phages have to favor lysis over lysogeny even though their progeny will most likely not be able to find any susceptible bacteria to infect. Even though the tradeoff can thus not be escaped by means of an *ENC*- dependent probability of lysogenization, sensing the number of coinfections could still attenuate it, in part, by utilizing the fact that the residual probability to belong to the early phase of the infection dynamics is approximately tenfold higher for infections by a single phage as compared to coinfections (Fig. 2A). Given that some phages have to favor lysis over lysogeny to allow for a quick initial spread, it should thus at least be possible to minimize their number—and thus the negative effect on the fraction of lysogenized bacteria—if predominantly infections with an *ENC* of one result in lysis.

To test this hypothesis, we analyzed how the infection dynamics of the *ENC*-sensitive phage variant change depending on the degree with which the probability of lysogenization depends on the *ENC*. We thereby assumed that the dependency of the probability of lysogenization on the *ENC* resembles a “switch” between a probability of zero for bacteria infected by a single phage, and some non-zero, variable probability *𝒫*_*ENC*≥2_ for all coinfections. We found a nearly linear relationship between the concentration of lysogens at the end of the infection dynamics and the value of *𝒫*_*ENC*≥2_ (Fig. 4A). In contrast, the onset of the critical phase of the infection dynamics was insensitive to the value of *𝒫*_*ENC*≥2_, and occurred at similar times as for *ENC*-insensitive phages with a constant probability of lysogenization of around 1% (Fig. 4A). This indicates that the phage can maximize the fraction of lysogenized bacteria without significantly delaying the take-over of the bacterial population by lysing all bacteria infected by a single phage and lysogenizing all coinfected bacteria. Beyond this point, a further increase in the fraction of lysogenized bacteria is only possible when also a certain fraction of infections with *ENC* of one would result in lysogeny, which would however come at the cost of significantly delaying the takeover of the bacterial population (Fig. 4A).

The switch-like fate decision algorithm described above, albeit simple, utilizes most of the information conveyed by the *ENC*, and thus can be considered close to optimal from an information theoretical point of view (*I(Phase; Fate*)/*I(Phase; ENC*) = 76% for the infection dynamics shown in Fig. 1C, with *Fate* ∈ {*lysis, lysogeny*}). Indeed, most of the information not utilized concerns the phage’s theoretical ability to also show distinct fate decision probabilities after the end of the critical phase, which would however not further attenuate the tradeoff between phage growth and the fraction of lysogenized bacteria (when not distinguishing between the critical phase and the phase thereafter, 97% of the information is utilized). Note, that the experimentally determined dependency of the probability of lysogenization on the *ENC* in phage λ (0.38%, 69.60% and 98.86% for an *ENC* of one, two, and more than two, respectively, see (26)) resembles the close-to-optimal, switch-like fate decision algorithm described above (Fig. 4A). This suggests that phage λ maximized the fraction of lysogenized bacteria under the condition that the take-over of the bacterial population must occur as quickly as possible.

To confirm that an *ENC*-dependent probability of lysogenization provides temperate phages with a competitive advantage, we simulated the dynamics of a bacterial population coinfected with equal initial numbers of *ENC*-sensitive and *ENC*-insensitive phages (Fig. 4B&C), and tracked the ratio between the two types of phages as the infection dynamics progressed. We set the probability of lysogenization of the *ENC*-insensitive competitor to 36%, such that an independent infection by either *ENC*-sensitive or *ENC*-insensitive phages would result in approximately the same final concentration of lysogens. Early during this scenario, the initially low ⟨*ENC*(*t*)⟩ resulted in a small growth advantage for the *ENC*-sensitive phage (Fig. 4B). This small advantage accumulated over time and, at around four hours into the infection dynamics, the concentration of *ENC*-sensitive phages was around three times greater than the concentration of *ENC*-insensitive ones (Fig. 4C). On the other hand, nearly all lysogens at this time carried the *ENC*-insensitive prophage (Fig. 4C). This changed dramatically at around four and a half hours, when the ⟨*ENC*(*t*)⟩ substantially increased and the concentration of lysogens carrying the *ENC*-sensitive prophage surpassed that of the *ENC*-insensitive prophage (Fig. 4C). Since, after the critical phase, the only possibility to produce significant numbers of free phages is the induction of lysogens, the ratio between the two phage types converged to the ratio between the lysogens that produced them (assuming equal induction probabilities). This suggests that the *ENC*- dependency of the fate decision provides temperate phages with a selective advantage, and we found that this advantage persisted regardless of the competitor’s probability of lysogenization, which underscores the robustness of our results (Fig. 4D).

Finally, we asked whether the competitive advantage of *ENC*-sensitive temperate phages is sufficient for these phages to increase in frequency and eventually fix in populations initially dominated by *ENC*-insensitive competitors. To this end, we started our simulations with an initial population of 1000 phages, of which only 1% were *ENC*- sensitive. We then repeated the simulation several times mimicking a serial transfer experiment, with the initial fraction of *ENC*-sensitive phages in a subsequent simulation corresponding to the final fraction at the end of the infection dynamics in the previous one. We found that, in this series of transfers, the fraction of *ENC*-sensitive phages increased in frequency and fixed after at most ten transfers, regardless of the probability of lysogenization of the *ENC*-insensitive competitor phage (Fig. 4E). This result supports the hypothesis that the ability of temperate phages to increase the probability of lysogenization in response to increasing *ENC*s provides temperate phages with a robust selective advantage.

## Discussion

Prophages are a universal feature of bacterial genomes (39), and many genomic studies point towards the importance of these genetic elements for bacterial ecology and evolution (40; 41; 42; 43; 44; 45). Since the 1950’s, the molecular biology of several phages has been intensively studied and is now relatively well understood (18; 22; 46). However, connecting the underlying molecular biology with genomic data obtained from sequencing studies is non-trivial and requires understanding of the basic interaction dynamics between phages and bacteria at the population level (11; 47). Here, we used a stochastic modeling approach to gain insight into how coinfection of bacteria by multiple phages, a process occurring at the level of individual bacterial cells, affects the overall phage-host population dynamics and vice versa.

Despite the limited information conveyed by coinfections to the individual phage about the state of the infection at the population level, we found that the ability of phages to adjust the probability of lysogenization in response to increased numbers of coinfections can lead to a faster and more efficient takeover of bacterial populations. This demonstrates that even the utilization of minimal sensory information can lead to vast fitness increases if the underlying process involves large amplification steps (in this case the exponential growth of the phage) (48; 49). To be effective, the respective regulatory mechanism however has to take maximal advantage of the very limited information available (49). Our result showing that the lysis of all bacteria infected by a single phage and the lysogenization of all coinfected bacteria maximizes the exploitation of the information conveyed by the *ENC*, might thus explain the experimentally observed sharp increase of the probability of lysogenization upon coinfection by multiple phages (26). In contrast, a more gradual dependency of the *ENC* on the number of coinfections would require increasing the probability of coinfections and thus the information conveyed by the *ENC* to individual phages. Assuming that phage growth is already maximized, Eq. 2 suggests that the only way to increase the number of coinfections would be by prolonging the fate decision period. Interestingly, the lytic pathway is the default mode of phage λ development, and the decision to switch to the lysogenic pathway occurs when the lytic pathway genes *O* and *P* are already expressed (18). In lysogenic development, the activity of these two genes involved in DNA replication is actively repressed (50), as phage DNA replication is lethal for lysogens (51). As compared to a fate decision preceding the expression of *O* and *P*, this molecular mechanism can thus be interpreted to already delay the fate decision and thus to maximize the *ENC*. This delay thus leads to an informational edge for the phage while still allowing for fast phage replication in case of lytic development.

Recently, switching of viral communities from lysis to lysogeny in natural environments with high microbial concentrations has been proposed (3) and debated (52). We have previously shown experimentally that postponing the onset of infection until higher bacterial concentrations in bacteria carrying innate immune systems can increase prophage acquisition rates and attributed this effect to cell-density-dependent physiological changes (11). Notably, the increased occurrence of confections at high bacterial concentrations described in the present work is independent of such physiological effects (Fig. 2D). Driven purely by the phage-host population dynamics, this effect thus represents an additional mechanism that can lead to an increase in the prophage acquisition rates at high bacterial concentrations. Both, the physiological and the population-dynamic effects might act together in natural populations and thus help explain the observed higher prevalence of prophages in denser bacterial populations across diverse environments (3).

Given the experimentally observed strong dependency of the probability of lysogenization on the number of coinfecting phages, our result that all fate decisions, even those based on a single phage infection, most likely occur during the critical phase of the infection dynamics comprising the last horizontal reproduction cycle of the phage, strongly suggests that phages do not solely base their fate decision on the anticipated availability of susceptible bacteria. This somewhat counter-intuitive result can be explained by the fact that the fate decisions of individual phages are not (strictly) independent, since only those phages which are released by lysing bacteria can decide on bacterial fates in the first place. This interdependency forces the fate decision of phages into a seemingly cooperative mode, whereby bacteria infected by only a single phage are lysed to allow for a quick initial spread of the phage population, even though, in the majority of such infections, the released phage progeny will not be able to find any susceptible bacteria to infect (Fig. 3A). Why do individual phages not “defect this common goal” by acquiring mutations favoring lysogeny? The potential advantages of belonging to the first phages infecting a population of susceptible bacteria, even though being unlikely, are tremendous in terms of spread of genetic information. Thus, favoring lysis over lysogeny might still on average be beneficial even if it is usually detrimental. Even if this is not the case, the prophages descending from “defecting phages” will have lost the ability to quickly take over subsequent populations of susceptible bacteria once induced, which corresponds to a significant long-term selective disadvantage of defecting.

We have shown that the factors determining the widespread occurrence of coinfections by multiple phages differ widely depending on whether the phage is inoculated at high phage concentrations, or if it has to reach these concentrations itself by successive cycles of horizontal reproduction (Box 1). However, like most theoretical studies, we still assumed that the phage-host interaction dynamics take place in a well-mixed environment. Nevertheless, we think that our results should still qualitatively hold for semi-structured bacterial populations where the probability of coinfections does not solely depend on the spatial proximity to the nearest lysing bacterium. However, one should be careful not to simply assume that this has to always be the case, or that our results could be directly generalized to highly structured environments like dense biofilms. At the very least, we think that our study can serve as a starting point for a thorough analysis of how the factors determining the probability of multiple coinfections change, depending on environments in which bacteria and phages interact.

## Materials and Methods

### Preliminaries

The infection dynamics of a population of susceptible bacteria by temperate phages are shaped both by processes of a stochastic nature like the random collisions of bacteria and phages (36), and processes which have rather uniform durations with low coefficients of variations like the time of bacterial lysis after infection (33; 38; 53; 54) (see (55) for a recent review). Consequently, our stochastic model is based on two different reaction types: “Propensity reactions”, which we denote by a simple arrow, are assumed to have exponentially distributed firing times. For example, the propensity reaction 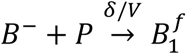 describes the attachment and subsequent infection of a susceptible bacterium *B*^−^ by a phage *P* with rate constant *δ*/*V*, resulting in an infected bacterium 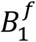 that is not yet committed to a given fate (lysis or lysogeny), where the subscript indicates that the number of phage coinfections is currently one. On the other hand, we describe processes having rather uniform durations with small coefficients of variation by “time-delay reactions”, which are denoted by a double arrow. For example, the time-delay reaction 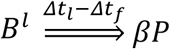 describes the lysis of a bacterium *B*^*l*^ at exactly *Δt*_*l*_ − *Δt*_*f*_ time units after it has committed to the lytic fate, resulting in the release of *β* copies of the phage *P*.

### Algorithmic Implementation

From a computational as well as conceptual point of view, the combination of propensity and time-delay reactions (see previous section) is challenging because it requires to record for each infected bacterium separately the number of phage coinfections as well as the time of its first infection. Conceptually, we accomplish this by applying a semi rule-based framework, while implementation-wise using a mixed stochastic/agent-based approach automatically minimizing the set of species for which element-wise information must be recorded, thus reducing the computational load. We implemented this framework in the open-source stochastic simulator *stochsim* freely available under https://langmo.github.io/stochsim/, which also includes all files necessary to simulate the model analyzed in this article. While *stochsim* is implemented in C++ to allow for the efficient simulation of the interactions between billions of bacteria and phages, it provides a convenient mini-language to configure a stochastic model, together with a MEX interface to MATLAB (The MathWorks, Inc., Natick, Massachusetts, United States) to configure, run and analyze the simulations. In the following, we conveniently use a simplified notation for the description of our model that does not fully reflect all details of the applied framework. However, we stress that the simulations were run such that the latent and the fate decision period always had a fixed length independently of the numbers of phage coinfections.

### Stochastic Model of Infection Dynamics

Our stochastic model describes the spread of temperate phages in a population of susceptible bacteria in a well-mixed habitat of volume *V*. It is based on and significantly extends our previous model described in (11). In this model, the species *P* denotes free phages. For the bacterial population, separate species denote susceptible but yet uninfected bacteria (*B*^−^), recently infected bacteria which did not yet commit to a fate (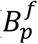, with *p* ≥ 1 the number of phage coinfections), bacteria committed to lysis which did not yet burst (*B*^*l*^), and lysogens (*B*^+^). We assume that phages attach to all bacteria with the same rate constant *δ*. Immediately after attachment, phages are assumed to adsorb and infect susceptible bacteria. Adsorption of phages to lysogens or to infected bacteria which have already committed to a given fate (lysis or lysogeny) is assumed to have no effect (12; 18). Infected bacteria are assumed to commit to a given fate after a fixed time period following their first infection, to which we refer as the fate decision period *Δt*_*f*_. They commit to the lytic fate with probability 1 − *𝒫*^*l*^(*p*), and to the lysogenic fate with probability *𝒫*^*l*^(*p*). Depending on the simulation, the probability of lysogenization is either assumed to be constant 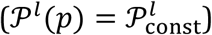, or to depend on the number of phage coinfections 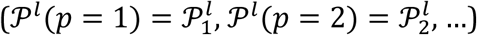. Bacteria committed to lysis are assumed to burst and to release *β* (burst size) phages *Δt*_*l*_ time units after their first infection, with *Δt*_*l*_ referred to as the latent period. Uninfected bacteria and lysogens are assumed to grow exponentially with the same doubling rate constant *v*_*d*_ (20), while bacteria committed to the lytic pathway are assumed not to divide (5 p. 26; 23). Finally, we assume that lysogens spontaneously enter the lytic pathway upon induction of the prophage (18) with the induction rate constant *ξ*.

Given these assumptions, our model is described by the following reactions:

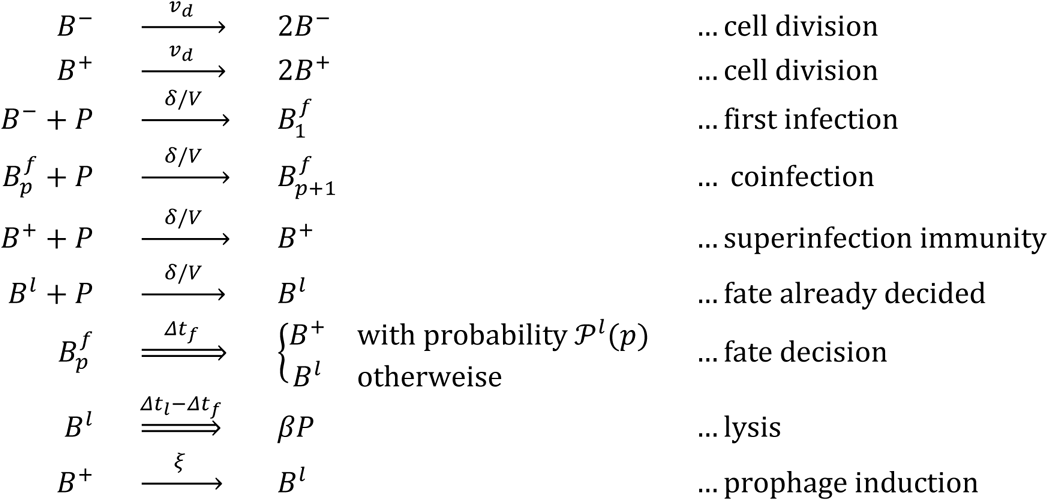

The model is initialized with zero concentrations for all species except for uninfected bacteria (*B*^−^) and free phages (*P*), for which the initial concentrations are indicated in the respective figure legends.

### Model for Coinfection by *ENC*-Sensitive and -Insensitive Phages

To simulate the coinfection of a bacterial population by *ENC*-sensitive (subscript *s*) and *ENC*-insensitive phages (subscript *i*), we modified our model to contain two separate species each for the free phages (*P*_*s*_ and *P*_*i*_), lysogens (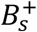 and 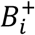) and bacteria committed to the lytic fate (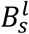 and 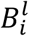), and replicated the corresponding reactions. For recently infected bacteria which did not yet commit to a given fate 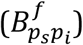, we counted the number of coinfections *p*_*s*_ ≥ 0 and *p*_*i*_ ≥ 0, *p*_*s*_ + *p*_*i*_ ≥ 1, by *ENC*-sensitive and *ENC*-insensitive phages separately. After the fate decision period, bacteria infected by either the *ENC*- sensitive or *ENC*-insensitive phage individually commit to a given fate as described above, while bacteria coinfected by both phage variants were assumed to commit to the lysogenic fate with a probability corresponding to the weighted average of the probabilities of the two phage variants:

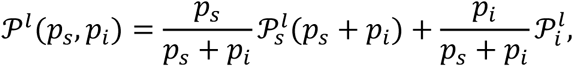

with 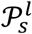 and 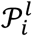 the probabilities of lysogenization of the *ENC*-sensitive and -insensitive phage, respectively. Once committed to a given fate, bacteria become lysogens of the *ENC*- sensitive or *ENC*-insensitive phage, respectively release these phages when bursting, with a probability proportional to the fraction of coinfections by the respective phage.

### Model Parametrization

Based on dedicated experiments and modelling, it was recently estimated that coinfections by bacteriophage λ occurring more than 24*min* after the first infection have negligible effect on the bacterial fate decision (27). However, already before this final threshold, the influence of subsequent coinfections on the bacterial fate decision decreases the later the respective phage arrives, with the influence of phages arriving between 5*min* and 15*min* after the first infection being only half as big as of phages arriving immediately after the first infection (estimated from Fig. 3B in (27), compare (33; 34)). Based on this experimental evidence, we assumed a fate decision period of *Δt*_*f*_ = 10*min* for our model if not indicated differently. Thus, coinfections are only considered for the calculation of the *ENC* if they occur up to *Δt*_*f*_ time units after the infection by the first phage.

For simulations where the probability of lysogenization depends on the number of phage coinfections *p*, we assumed a probability of lysogenization of *𝒫*_1_ = 0.0038 for one infection, of *𝒫*_2_ = 0.6960 for two coinfections, and of *𝒫*_*p*_ = 0.9886 for *p* ≥ 3 coinfections (26). For simulations assuming a constant probability of lysogenization, this probability was set to *𝒫*_const_ = 0.02 if not indicated differently. All other parameters were taken from our previous model (11). The complete set of parameters is given in Table 1.

## Acknowledgements

We are grateful to S. Abedon, C. Igler, M. Lagator, G. Tkačik, and P. Payne for comments on the manuscript. The research leading to these results has received funding from the People Programme (Marie Curie Actions) of the European Union’s Seventh Framework Programme (FP7/2007–2013) under REA Grant Agreement No. 291734 (ML). MP is a Simons foundation fellow of the Life Sciences Research Foundation. This work was funded by an HFSP Young Investigators’ grant (CCG).

## Bibliography

1. Clokie, MRJ, et al. Phages in nature. Bacteriophage. 2011, Vol. 1, 1, pp. 31–45.

2. Bergh, Ø, et al. High abundance of viruses found in aquatic environments. Nature. 1989, Vol. 340, pp. 467–468.

3. Knowles, B, et al. Lytic to temperate switching of viral communities. Nature. 2016, Vol. 531, pp. 466–470.

4. Levin, BR, Stewart, FM and Chao, L. Resource-limited growth, competition, and predation: a model and experimental studies with bacteria and bacteriophage. The American Naturalist. 1977, Vol. 111, pp. 3–24.

5. Weitz, JS. Quantitative viral ecology: dynamics of viruses and their microbial hosts. Princeton : Princeton University Press, 2015. Vol. 55.

6. Lenski, RE and Levin, BR. Constraints on the coevolution of bacteria and virulent phage: a model, some experiments, and predictions for natural communities. The American Naturalist. 1985, Vol. 125, pp. 585–602.

7. Westra, ER, et al. Parasite exposure drives selective evolution of constitutive versus inducible defense. Current Biology. 2015, Vol. 25, pp. 1043–1049.

8. Chao, L, Levin, BR and Stewart, FM. A complex community in a simple habitat: an experimental study with bacteria and phage. Ecology. 1977, Vol. 58, pp. 369–378.

9. Buckling, A and Rainey, PB. Antagonistic coevolution between a bacterium and a bacteriophage. Proceedings of the Royal Society of London B: Biological Sciences. 2002, Vol. 269, pp. 931–936.

10. Gómez, P and Buckling, A. Bacteria-phage antagonistic coevolution in soil. Science. 2011, Vol. 332, pp. 106–109.

11. Pleška, M, et al. Phage-host population dynamics promotes prophage acquisition in bacteria with innate immunity. Nature ecology & evolution. 2018, p. 1.

12. Stewart, FM and Levin, BR. The population biology of bacterial viruses: why be temperate. Theoretical population biology. 1984, Vol. 26, pp. 93–117.

13. Bossi, L, et al. Prophage contribution to bacterial population dynamics. Journal of bacteriology. 2003, Vol. 185, pp. 6467–6471.

14. -Gama, JA, et al. Temperate bacterial viruses as double-edged swords in bacterial warfare. PLoS One. 2013, Vol. 8, p. e59043.

15. Erez, Z, et al. Communication between viruses guides lysis-lysogeny decisions. Nature. 2017, Vol. 541, pp. 488–493.

16. Payne, RJH and Jansen, VAA. Understanding bacteriophage therapy as a density-dependent kinetic process. Journal of Theoretical Biology. 2001, Vol. 208, 1.

17. Kourilsky, P. Lysogenization by bacteriophage lambda: II.-identification of genes involved in the multiplicity dependent processes. Biochimie. 1975, Vol. 56, pp. 1511–1516.

18. Oppenheim, AB, et al. Switches in bacteriophage lambda development. Annu. Rev. Genet. 2005, Vol. 39, pp. 409–429.

19. Howard-Varona, C, et al. Lysogeny in nature: mechanisms, impact and ecology of temperate phages. The ISME journal. 2017, Vol. 11, 7, pp. 1511–1520.

20. Lieb, M. The establishment of lysogenicity in Escherichia coli. Journal of bacteriology. 1953, Vol. 65, p. 642.

21. Lwoff, A. Lysogeny. Bacteriological Reviews. 1953, Vol. 17, p. 269.

22. Ptashne, M. A genetic switch: Gene control and phage Lambda. s.l. : Palo Alto, CA (US); Blackwell Scientific Publications, 1986.

23. St-Pierre, F and Endy, D. Determination of cell fate selection during phage lambda infection. Proceedings of the National Academy of Sciences. 2008, Vol. 105, pp. 20705–20710.

24. Kourilsky, P. Lysogenization by bacteriophage lambda. I. Multiple Infection and the Lysogenic Response. Molecular and General Genetics. 1973, Vol. 122, pp. 183–195.

25. Zeng, L, et al. Decision making at a subcellular level determines the outcome of bacteriophage infection. Cell. 2010, Vol. 141, pp. 682–691.

26. Avlund, M, et al. Why Do Phage Play Dice? Journal of virology. 2009, Vol. 83, pp. 11416–11420.

27. Cortes, MG, et al. Late-Arriving Signals Contribute Less to Cell-Fate Decisions. Biophysical Journal. 2017, Vol. 113, pp. 2110–2120.

28. Fry, BA. Conditions for the infection of Escherichia coli with lambda phage and for the establishment of lysogeny. Microbiology. 1959, Vol. 21, pp. 676–684.

29. Levine, M. Mutations in the temperate phage P22 and lysogeny in Salmonella. Virology. 1957, Vol. 3, pp. 22–41.

30. Bertani, LE. The effect of the inhibition of protein synthesis on the establishment of lysogeny. Virology. 1957, Vol. 4, pp. 53–71.

31. Kourilsky, P and Knapp, A. Lysogenization by bacteriophage lambda: III.-multiplicity dependent phenomena occuring upon infection by lambda. Biochimie. 1975, Vol. 56, pp. 1517–1523.

32. Kourilsky, P and Gros, D. Lysogenization by bacteriophage lambda IV inhibition of phage DNA synthesis by the products of genes cII and cIII. Biochimie. 1977, Vol. 58, pp. 1321–1327.

33. Singh, A and Dennehy, JJ. Stochastic holin expression can account for lysis time variation in the bacteriophage λ. Journal of The Royal Society Interface. 2014, Vol. 11, p. 20140140.

34. Weitz, JS, et al. Collective decision making in bacterial viruses. Biophysical journal. 2008, Vol. 95, pp. 2673–2680.

35. Payne, RJH and Jansen, VAA. Phage therapy: the peculiar kinetics of self-replicating pharmaceuticals. Clinical pharmacology & therapeutics. 2000, Vol. 68, 3.

36. Kasman, LM, et al. Overcoming the phage replication threshold: a mathematical model with implications for phage therapy. Journal of virology. 2002, Vol. 76, pp. 5557–5564.

37. Ellis, EL and Delbrück, M. The growth of bacteriophage. The Journal of general physiology. 1939, Vol. 22, pp. 365–384.

38. Dennehy, JJ and Wang, N. Factors influencing lysis time stochasticity in bacteriophage λ. BMC microbiology. 2011, Vol. 11, p. 174.

39. Canchaya, C, et al. Prophage genomics. Microbiology and Molecular Biology Reviews. 2003, Vol. 67, pp. 238–276.

40. Obeng, N, Pratama, AA and Elsas, JD. The significance of mutualistic phages for bacterial ecology and evolution. Trends in microbiology. 2016, Vol. 24, pp. 440–449.

41. Brüssow, H, Canchaya, C and Hardt, W-D. Phages and the evolution of bacterial pathogens: from genomic rearrangements to lysogenic conversion. Microbiology and molecular biology reviews. 2004, Vol. 68, pp. 560–602.

42. Casjens, S. Prophages and bacterial genomics: what have we learned so far? Molecular microbiology. 2003, Vol. 49, pp. 277–300.

43. Bobay, L-M, Touchon, M and Rocha, EPC. Manipulating or superseding host recombination functions: a dilemma that shapes phage evolvability. PLoS genetics. 2013, Vol. 9, p. e1003825.

44. Bobay, L-M. Pervasive domestication of defective prophages by bacteria. Proceedings of the National Academy of Sciences. 2014, Vol. 111, pp. 12127–12132.

45. Touchon, M, Moura de Sousa, JA and Rocha, EPC. Embracing the enemy: the diversification of microbial gene repertoires by phage-mediated horizontal gene transfer. Current opinion in microbiology. 2017, Vol. 38, pp. 66–73.

46. Bertani, G. Lysogeny at mid-twentieth century: P1, P2, and other experimental systems. Journal of bacteriology. 2004, Vol. 186, pp. 595–600.

47. Mitarai, N, Brown, S and Sneppen, K. Population dynamics of phage and bacteria in spatially structured habitats using Phage λ and Escherichia coli. Journal of bacteriology. 2016, Vol. 198, pp. 1783–1793.

48. Tkačik, G and Bialek, W. Information processing in living systems. Annual Review of Condensed Matter Physics. 7, 2016, pp. 89–117 .

49. Taylor, SF, Tishby, N und Bialek, W. Information and fitness. arXiv preprint 0712.4382. 2007.

50. Mensa-Wilmot, K, Carroll, K und McMacken, R. Transcriptional activation of bacteriophage lambda DNA replication in vitro: regulatory role of histone-like protein HU of Escherichia coli. The EMBO Journal. 1989, Bd. 8, 8, S. 2393–2402.

51. Brachet, Philippe, Eisen, Harvey and Rambach, Alain. Mutations of coliphage lambda affecting the expression of replicative functions O and P. Molecular and General Genetics. 1970, Vol. 108, 3, pp. 266–276.

52. Weitz, JS, et al. Lysis, lysogeny and virus-microbe ratios. Nature. 2017, Vol. 549, p. E1.

53. Amir, A, et al. Noise in timing and precision of gene activities in a genetic cascade. Molecular Systems Biology. 2007, Vol. 3, p. 71.

54. Wang, I-N, Smith, DL and Young, R. Holins: the protein clocks of bacteriophage infections. Annual Reviews in Microbiology. 2000, Vol. 54, pp. 799–825.

55. Golding, I. Infection by bacteriophage lambda: an evolving paradigm for cellular individuality. Current Opinion in Microbiology. 2018, Vol. 43, pp. 9–13.

56. Abedon, ST. Phage therapy dosing: The problem(s) with multiplicity of infection (MOI). Bacteriophage. 2016, Vol. 6, 3, p. e1220348.

57. Goodwin, GC, Graebe, SF and Salgado, ME. Control System Design. Englewood Cliffs : Prentice Hall, 2001.

58. Abedon, ST. Commentary: Communication between Viruses Guides Lysis-Lysogeny Decisions. Frontiers in Microbiology. 2017, Vol. 8.

59. Abedon, ST. Selection for lysis inhibition in bacteriophage. Journal of theoretical biology. 1990, Vol. 146, pp. 501–511.

60. Abedon, ST. Thinking about microcolonies as phage targets. Bacteriophage. 2012, Vol. 2, pp. 200–204.

61. Chibani-Chennoufi, S, et al. Phage-Host Interaction: an Ecological Perspective. Journal of Bacteriology. 2004, Vol. 186, 12.

62. Gillespie, DT. Exact stochastic simulation of coupled chemical reactions. The Journal of Physical Chemistry. 1977, Vol. 81, pp. 2340–2361.

63. Sutherland, William J, et al. Identification of 100 fundamental ecological questions. Journal of ecology. 2013, Vol. 101, 1.

64. Gandon, S. Why Be Temperate: Lessons from Bacteriophage λ. Trends in microbiology. 2016, Vol. 24, 5.

