## Supporting Information text and figures for "Population dynamics of decision making in temperate bacteriophages"

Moritz Lang, Maroš Pleška and Călin C. Guet

#### Supporting Information Text

In the following, we derive the analytical results stated in the main text. In our derivations, we approximate the stochastic dynamics (except the ones concerning coinfections) by their respective deterministic counterparts, which is valid given sufficiently large volumes.

##### Expected *ENC* at time *t*

In [4, p.49ff], it was derived that, in a well-mixed environment, the probability of a bacterium being infected by a phage per unit time only depends on the phage concentration. Since this also holds for bacteria already infected by a phage but not yet committed to a given fate (lysis or lysogeny), we can calculate the average number of phage coinfections (excluding the initial one) occurring during the fate decision period $\Delta t_f$ of a bacterium infected by its first phage at time $t$:

$$\begin{aligned}\langle ENC(t)\rangle - 1 =& \int_0^{\Delta t_f} \delta P(t+\tau)d\tau \\ =&\delta P(t)\int_0^{\Delta t_f} e^{v_{rel}(t+\tau)\tau}d\tau\end{aligned}$$

where $v_{rel}(t) := \frac{d}{dt}\log(P(t)) = \frac{\frac{d}{dt}P(t)}{P(t)}$ denotes the relative growth rate of the phage at time $t$. Since the relative growth rate $v_{rel}(t)$ of the phage changes only slowly during the relevant phases of the infection, the average $ENC$ at time $t$ becomes

$$\langle ENC(t)\rangle =1 + c(t)\delta P(t)\Delta t_f,$$

with

$$\begin{aligned}c(t) =& \frac{e^{v_{rel}(t)\Delta t_f} - 1}{v_{rel}(t)\Delta t_f} \\ \approx& 1 + \frac{1}{2}v_{rel}(t)\Delta t_f\end{aligned}$$

a factor taking into account that, already during the course of a relatively short fate decision period, the phage concentration can increase for a significant amount.

##### Average *ENC* during an infection dynamics

For the derivation of an analytical formula for the average $ENC$ during the whole infection dynamics, $\langle ENC\rangle_{tot}$, we assume that (nearly) all susceptible bacteria get eventually infected after some time $T$, that is, that the phage/host interaction dynamics are qualitatively similar to the ones shown in Figure 1C in the main text. The $\langle ENC\rangle_{tot}$ is the average of the $\langle ENC(t)\rangle$ weighted by the normalized rate $r_I(t) = \frac{\delta P(t)B^-(t)}{\int_0^T \delta P(t)B^-(t)dt}$ of infections of susceptible bacteria $B^-(t)$ by their first phage (see Box 1 in the main text):

$$\langle ENC\rangle_{tot} = \int_0^T \langle ENC(t)\rangle r_I(t)dt$$

$$\approx 1 + \delta \Delta t_f \frac{\int_0^T c(t) P^2(t) B^-(t) dt}{\int_0^T P(t) B^-(t) dt}.$$

Note that, mathematically, $\langle ENC \rangle_{tot}$ corresponds to the (unconditioned) expected value of the $ENC$ for an infected bacterium, $\langle ENC(t) \rangle$ to the expected $ENC$ conditioned on that the bacterium was infected by its first phage at time $t$, and $r_I(t)$ to the probability that a bacterium got infected by its first phage at time $t$, i.e.

$$\mathrm{E}[ENC] = \int_0^T \mathrm{E}[ENC | \text{first infection at } t] \, \mathrm{P}[\text{first infection at } t] dt.$$

Multiplying the differential equation for the concentration of uninfected bacteria on both sides by the product of the phage concentration $P(t)$ and the factor $c(t)$, and integrating (by parts) over time, we obtain

$$\begin{aligned} \frac{d}{dt} B^-(t) =& v_d B^-(t) - \delta P(t) B^-(t) \\ \Rightarrow \qquad \int_0^T c(t) P^2(t) B^-(t) dt =& \frac{v_d}{\delta} \int_0^T c(t) P(t) B^-(t) dt - \frac{1}{\delta} \int_0^T c(t) P(t) \frac{d}{dt} B^-(t) dt \\ =& \frac{v_d}{\delta} \int_0^T c(t) P(t) B^-(t) dt + \frac{1}{\delta} \int_0^T c(t) v_{rel}(t) P(t) B^-(t) dt \\ &+ \frac{1}{\delta} \int_0^T \frac{d}{dt} c(t) P(t) B^-(t) dt - \frac{1}{\delta} \left[ c(t) P(t) B^-(t) \right]_0^T. \end{aligned}$$

Inserting this partial solution into the equation for the average $ENC$ over the whole infection dynamics, we obtain

$$\langle ENC \rangle_{tot} \approx 1 + v_d \Delta t_f \langle c \rangle + \Delta t_f \langle c v_{rel} \rangle + \Delta t_f \langle c' \rangle - \Delta t_f \frac{\left[ c(t) P(t) B^-(t) \right]_0^T}{\int_0^T B^-(t) P(t) dt},$$

with $\langle c \rangle = \int_0^T c(t) r_I(t) dt$, $\langle c v_{rel} \rangle = \int_0^T c(t) v_{rel}(t) r_I(t) dt$ and $\langle c' \rangle = \int_0^T \frac{d}{dt} c(t) r_I(t) dt$ the weighted averages of the factor $c(t)$, of the product of $c(t)$ and the relative growth rate $v_{rel}(t)$, and of the rate of change $\frac{d}{dt} c(t)$ of the factor $c(t)$, respectively. For small initial phage concentrations $P(0)$ and sufficiently large $T$ such that (nearly) all susceptible bacteria are infected at the end of the infection dynamics (i.e. small $B^-(T)$), the last term in this sum is approximately zero, i.e.

$$\Delta t_f \frac{\left[ c(t) P(t) B^-(t) \right]_0^T}{\int_0^T B^-(t) P(t) dt} \approx 0,$$

and can thus be neglected.

By definition, 90% of all bacteria are infected by their first phage during the critical phase of the infection dynamics (see main text). Due to the averaging by the rate of new infections $r_I(t)$, thus also the values of $\langle c \rangle$, $\langle c v_{rel} \rangle$ and $\langle c' \rangle$ are nearly exclusively determined during the critical phase. Because the critical phase is rather short and because changes in the concentration of yet uninfected bacteria only impact phage growth with a delay given by the latent period, the relative growth rate of the phage changes only little during the critical phase. This implies that that $\langle c' \rangle \approx 0$, and that we can approximate $\langle c \rangle$ and $\langle c v_{rel} \rangle$ by their respective values at the critical infection time $t_{crit}$. We then obtain

$$\begin{aligned} \langle ENC \rangle_{tot} \approx& 1 + c(t_{crit}) v_{rel}(t_{crit}) \Delta t_f \left( 1 + \frac{v_d}{v_{rel}(t_{crit})} \right) \\ =& 1 + \left( e^{v_{rel}(t_{crit}) \Delta t_f} - 1 \right) \left( 1 + \frac{v_d}{v_{rel}(t_{crit})} \right). \end{aligned} \qquad (1)$$

If $v_{rel}(t_{crit}) \Delta t_f$ is sufficiently small, we can approximate the exponential by its first-order Taylor series approximation, and obtain

$$\langle ENC \rangle_{tot} \approx 1 + \Delta t_f (v_{rel}(t_{crit}) + v_d).$$

This final approximation however becomes less precise for high relative phage growth rates $v_{rel}(t_{crit})$, long fate decision periods $\Delta t_f$, or combinations thereof. In such cases, it is thus more accurate to use Eq. 1.

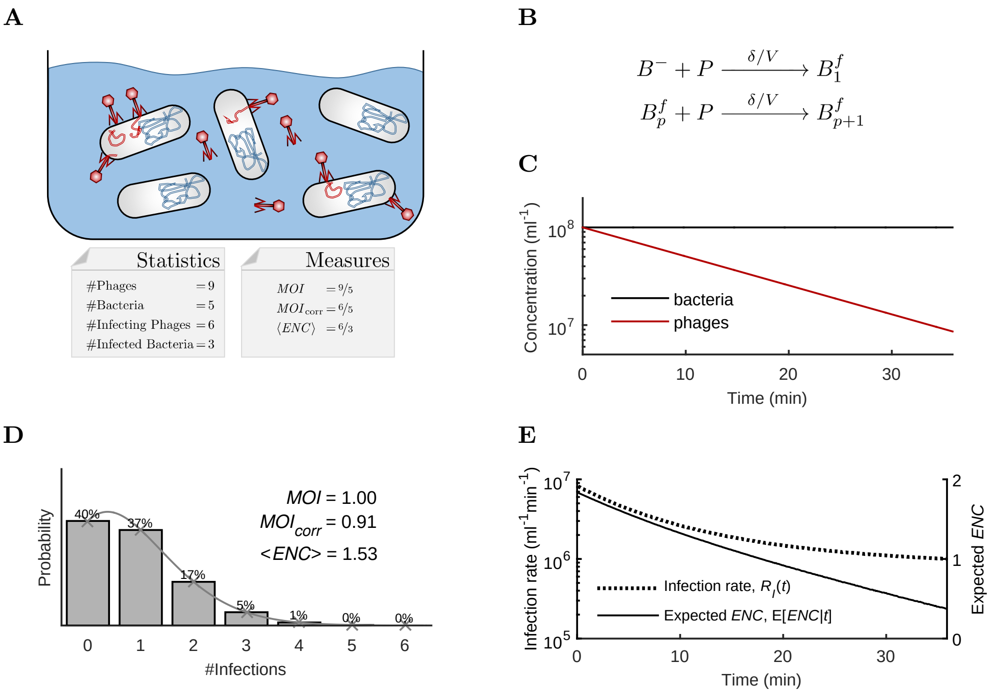


Supplementary Figure 1: Relationship between the *MOI* and the *ENC* in specialized phage assays in which phage replication, bacterial division, lysis, lysogeny, and the effect of different phage arrival times can be neglected. A: In the assay defined in [3], phages attach to bacteria under low temperatures, which slows down metabolism and prevents the insertion of the phage DNA into bacteria. After a sufficiently long time $T$ to allow most phages to attach to bacteria, the temperature is quickly increased such that nearly all phages insert their DNA into host bacteria at the same time. In such an assay, the *MOI* corresponds to the initial phage/bacterial ratio, and the average *ENC* to the ratio between the number of phages which infected bacteria and the number of bacteria infected by at least one phage. In [2], it was proposed to only take into account those phages which actually infected bacteria for the calculation of the *MOI* (referred to as $MOI_{\text{corr}}$) in agreement with the original definition [1]. B: Simplified version of our stochastic model for specialized phage assays as described in (A). The species and parameters correspond to those of our full model as described in the Materials and Methods. C: Dynamics of the simplified model for specialized phage assays. The bacterial concentration stays constant, while the concentration of free phages decreases exponentially. D: Distribution of the number of phage coinfections per cell encountered by susceptible bacteria for the infection dynamics shown in (C), and fit of a Poisson distribution with a mean equal to $MOI_{\text{corr}}$. Different to Figure 1E in the main text, also bacteria not infected by any phage were considered for this distribution in order to illustrate the Poisson fit. However, note that for the calculation of the average *ENC*, only bacteria infected by at least one phage are taken into account. For such specialized phage assays (and only then), $MOI_{\text{corr}} = (1 - \exp(-\delta B^- T))MOI$ [2], and $\langle ENC \rangle_{tot} = \frac{MOI_{\text{corr}}}{1-\exp(-MOI_{\text{corr}})}$. E: In contrast to the dynamic infection scenario discussed in the main text, the rate at which phages infect susceptible bacteria (solid curve, left axis) as well as the average *ENC* (dotted curve, right axis) of a bacterium infected at time $t$ decrease monotonically over time.

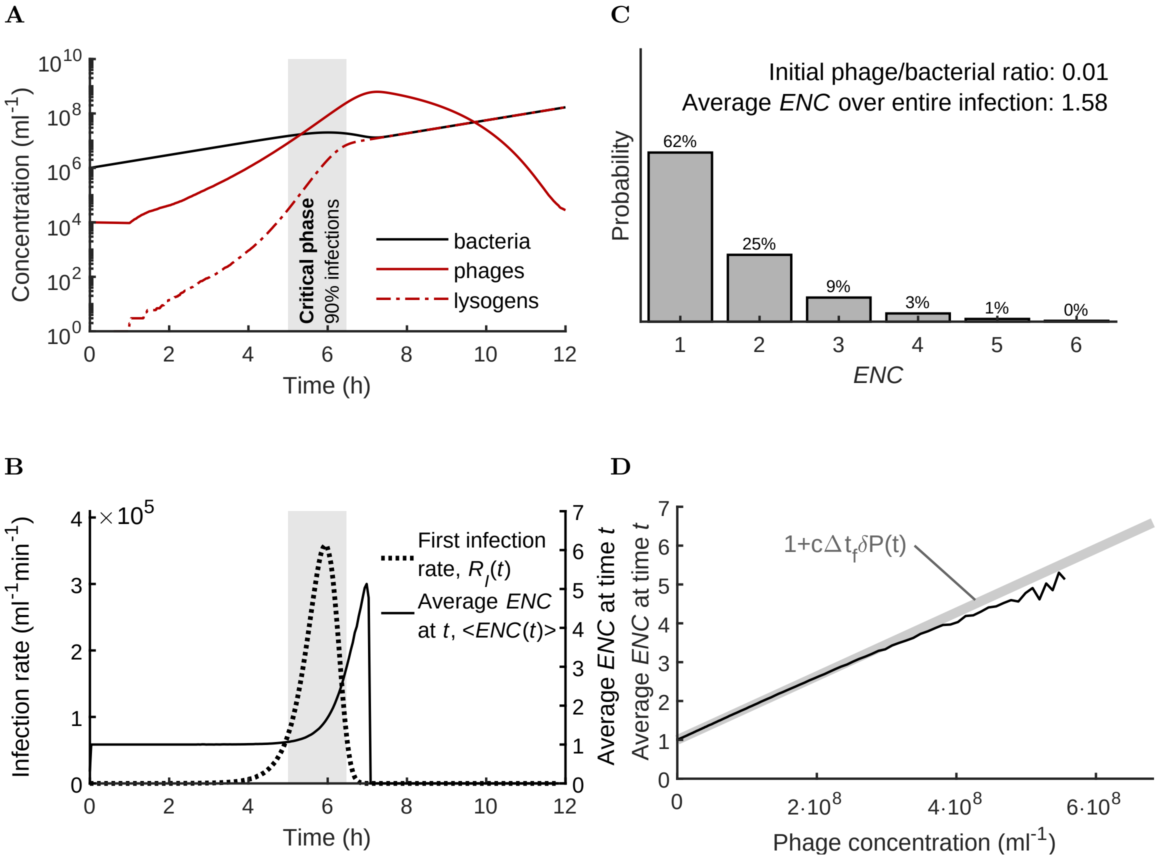


Supplementary Figure 2: Dynamics of a bacterial population infected by temperate phages. The dynamics correspond to the ones in Figure 1C-F in the main text when assuming an $ENC$-sensitive phage variant instead of an $ENC$-insensitive one. The probability of lysogenization of this variant was set to 0.38%, 69.60% and 98.86% for an ENC of one, two and more than two, respectively. All other details are identical to the ones described in Figure 1C-F in the main text.

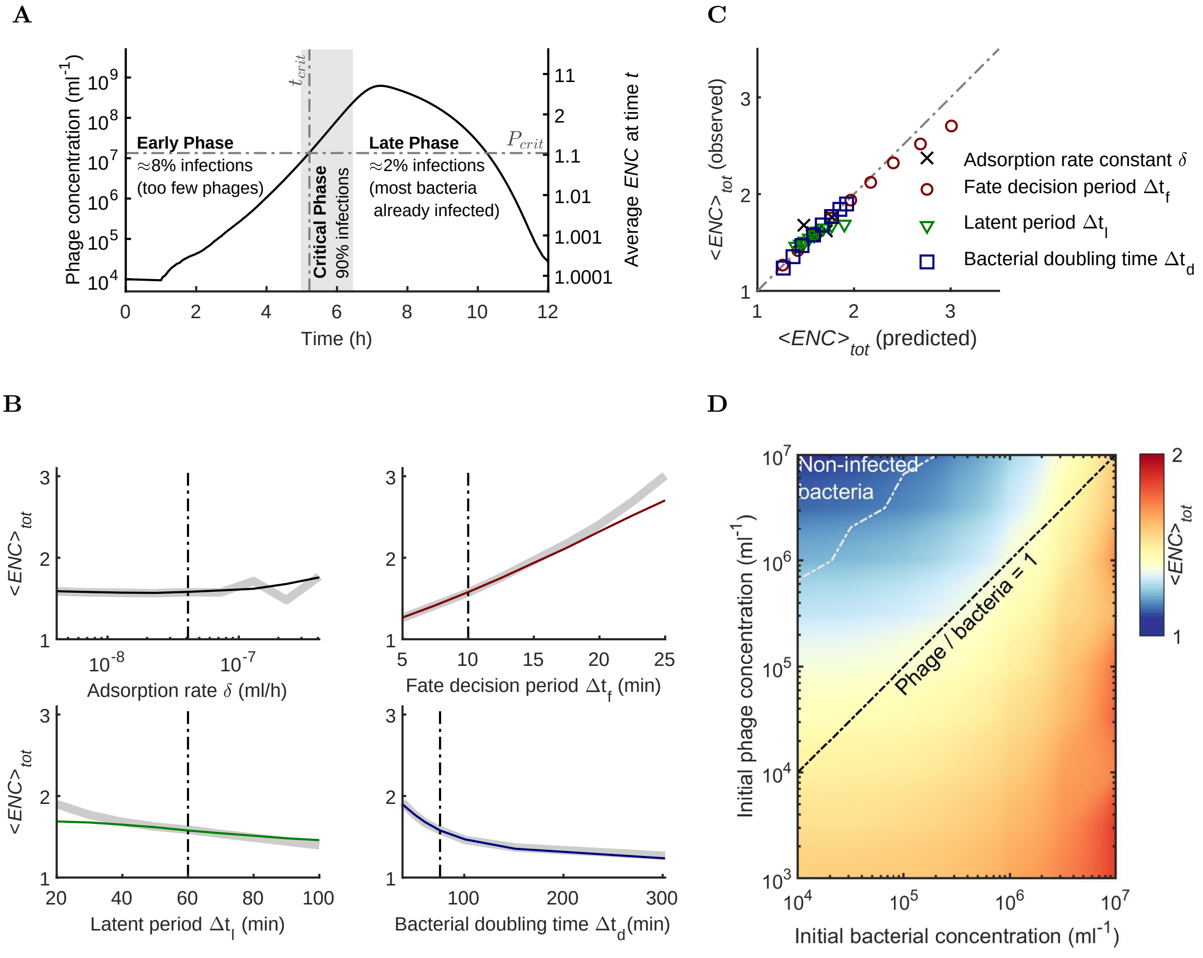


Supplementary Figure 3: Dependency of the ENC on key ecological and evolutionary parameters. The analysis in this figure corresponds to the one in Figure 2 in the main text when assuming an *ENC*-sensitive phage variant instead of an *ENC*-insensitive one. The probability of lysogenization of this variant was set to 0.38%, 69.60% and 98.86% for an ENC of one, two and more than two, respectively. All other details are identical to the ones described in Figure 2 in the main text.

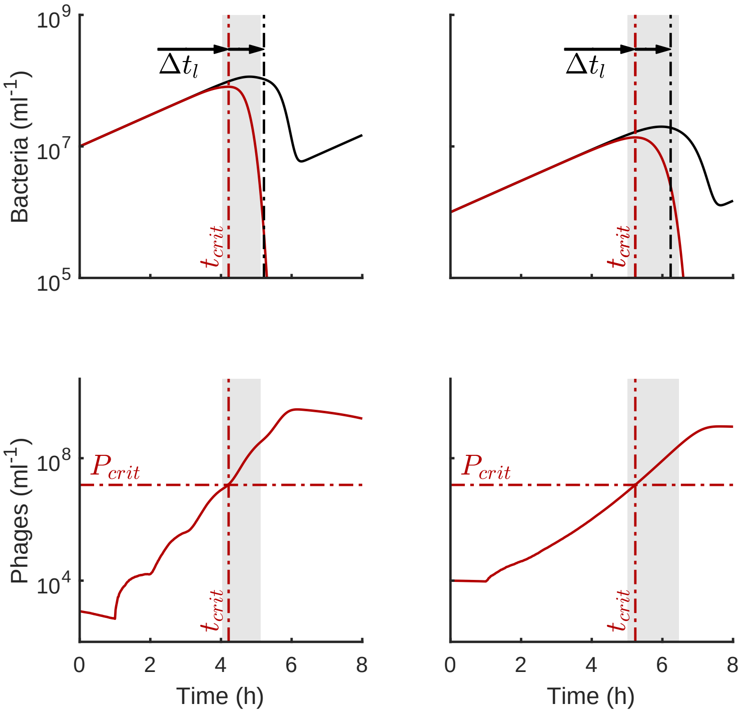


Supplementary Figure 4: Relationship between the critical phase and the critical phage threshold. The figures at the top show the dynamics of the total concentration of bacteria (black curve) and the ones of only those susceptible bacteria which were not yet infected (red). The figures at the bottom show the dynamics of the phage concentration. The initial phage and bacterial concentrations were set to $10^7 ml^{-1}$ and $10^3 ml^{-1}$ (left figures), respectively $10^6 ml^{-1}$ and $10^4 ml^{-1}$ (right figures). In both cases, the critical phase (light gray background) starts approximately at the time $t_{crit}$ (vertical red dash-dotted lines) when the phage reaches the critical concentration threshold $P_{crit}$ (horizontal red dash-dotted lines on bottom figures). This threshold $P_{crit}$ is independent of the initial conditions, and is thus the same in the left and the right column. At $t_{crit}$, the concentration of uninfected bacteria (red curves in top figures) stops to increase and starts to decrease. The total bacterial concentration (black curves), which also includes already infected bacteria and lysogens, however starts to decrease *after* $t_{crit}$, more specifically with a delay which has approximately the length of the latent period.

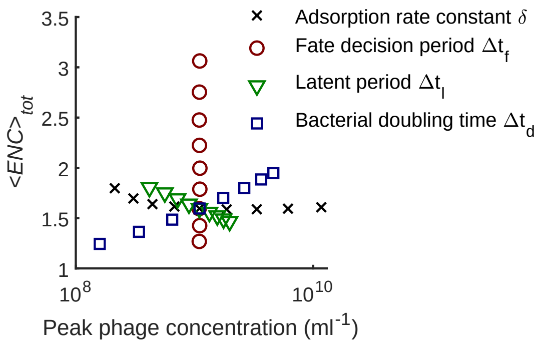


Supplementary Figure 5: Comparison between the average *ENC* over the whole infection dynamics obtained from the simulations in Figure 2B in the main text, and the peak phage concentration reached during the respective infection dynamics. Black crosses corresponding to varying the adsorption rate constant, red circles to varying the fate decision period, green triangles to varying the latent period, and blue squares to varying the bacterial doubling time.

### References

[1] Stephen T Abedon. Phage therapy dosing: The problem(s) with multiplicity of infection (MOI). *Bacteriophage*, 6(3):e1220348, 2016.

[2] Laura M Kasman, Alex Kasman, Caroline Westwater, Joseph Dolan, Michael G Schmidt, and James S Norris. Overcoming the phage replication threshold: A mathematical model with implications for phage therapy. *Journal of virology*, 76(11):5557–5564, 2002.

[3] Philippe Kourilsky. Lysogenization by bacteriophage lambda. I. Multiple infection and the lysogenic response. *Molecular and General Genetics*, 122(2):183–195, 1973.

[4] Joshua S Weitz, Yuriy Mileyko, Richard I Joh, and Eberhard O Voit. Collective decision making in bacterial viruses. *Biophysical journal*, 95(6):2673–2680, 2008.
